# Reprogramming protein interfaces between adenylation and carrier protein domains in nonribosomal peptide synthetases

**DOI:** 10.64898/2026.07.20.739675

**Authors:** Fumihiro Ishikawa, Iyo Arata, Taisei Uehara, Kana Kinoshita, Yuka Nakanishi, Kosei Sone, Toma Kashima, Fumitaka Kudo, Tadashi Eguchi, Tohru Terada, Shinya Fushinobu, Genzoh Tanabe, Akimasa Miyanaga

## Abstract

Nonribosomal peptide synthetases (NRPSs) assemble structurally diverse bioactive natural products through selective communication between adenylation (A) and carrier protein (CP) domains. Although rational rewiring of these protein–protein interactions could enable customized NRPS design, this remains challenging due to the dynamic nature of protein interfaces. Here we show that structure-guided interface reprogramming enables productive non-cognate A– CP pairings across enterobactin, vibriobactin, pyochelin, and vicenistatin biosynthetic systems. Engineered interactions between the A domains EntE, VibE, or PchD and the non-cognate CPs VinL or EntB were validated by biochemical, kinetic, and structural analyses. Reconstituted pathways containing engineered VibE, EntB, and EntF restored enterobactin production and increased yield to 2.2-fold that of the native EntE–EntB–EntF system. Interface-engineered PchD also enhanced production of a non-native salicylic acid–norspermidine conjugate. An X-ray structure and molecular dynamics simulations of an engineered non-cognate A–CP complex reveal recognition principles for programmable NRPS interface design.

## Introduction

Reprogramming of microbial biosynthetic assembly lines is a powerful strategy for sustainable access to structurally complex molecules with important medicinal properties. Nonribosomal peptides (NRPs) are particularly valuable resources for drug discovery due to their broad antibiotic, anticancer, and immunosuppressive activities. NRPs are biosynthesized by large, multifunctional enzymes called nonribosomal peptide synthetases (NRPSs).^1^ Adenylation (A) and carrier protein (CP) domains are common to all NRPS systems.^2^ A domains select and activate amino acid or aryl acid substrates as aminoacyl- or acyl-adenylate intermediates, using ATP, and then transfer the acyl group to the 4′-phosphopantetheine (Ppant) thiol of the CP domain to generate acyl-*S*-CP intermediates (Fig. 1A). Because A-domain substrates include proteinogenic and nonproteinogenic amino acids, keto acids, and aryl acids, A domains act as gatekeepers that control substrate incorporation into NRP products.^3^

**Fig. 1.**
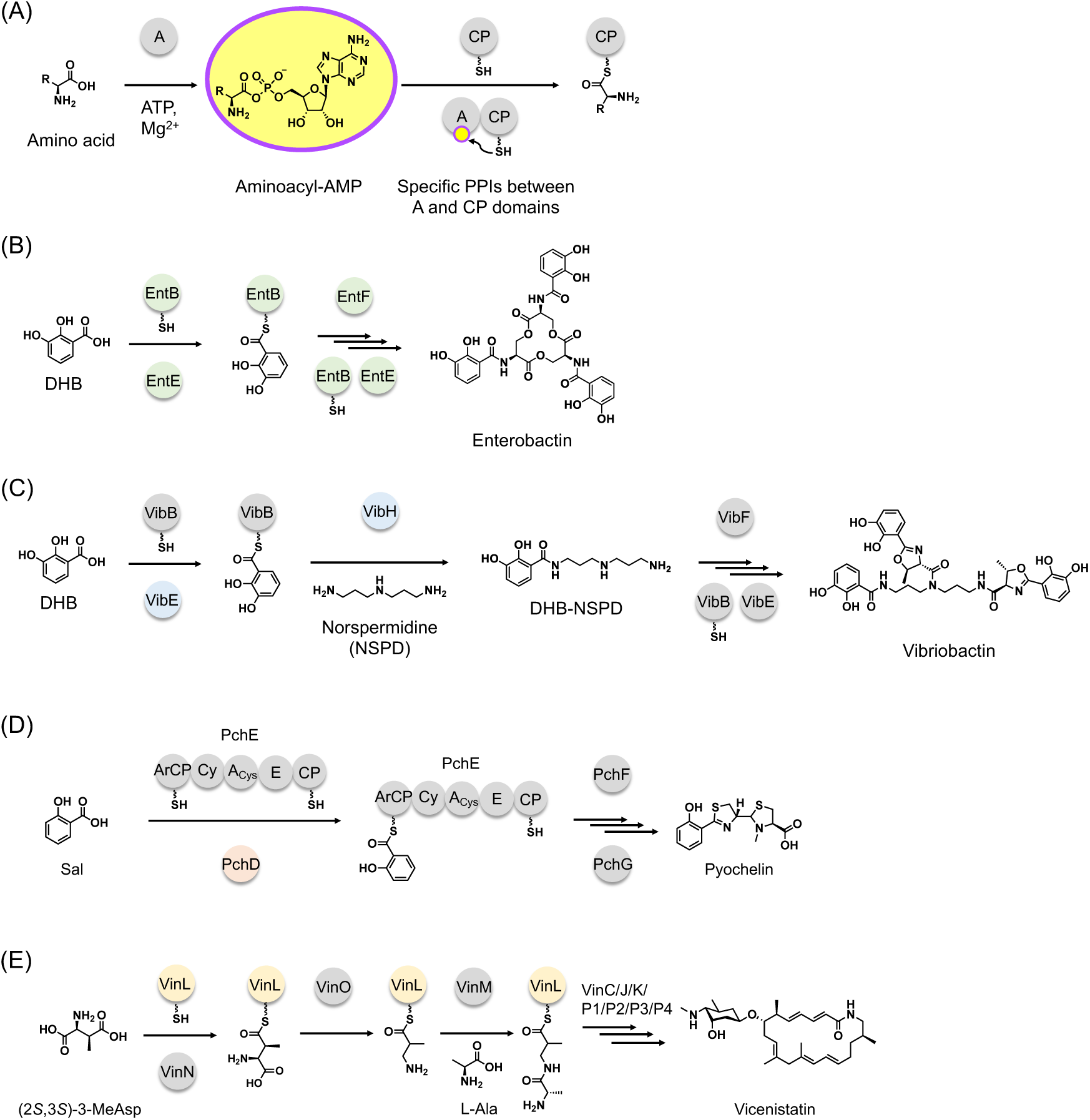
Substrate loading onto nonribosomal peptide synthetase (NRPS) assembly lines and the NRPS biosynthetic pathways described in this study. (A) Substrate activation and transfer catalyzed by adenylation (A) domains. Specific protein–protein interactions between A and carrier protein (CP) domains govern binding and, consequently, substrate loading. (B) Biosynthetic pathway of the siderophore enterobactin. The A domain EntE catalyzes the activation of 2,3-dihydroxybenzoic acid (DHB) and its transfer onto the aryl carrier protein (ArCP) domain of EntB. A detailed biosynthetic pathway of enterobactin is provided in Extended Data Fig. 2. (C) Biosynthetic pathway of the siderophore vibriobactin. The A domain VibE catalyzes the activation of DHB and its transfer onto the ArCP domain of VibB. (D) Biosynthetic pathway of the siderophore pyochelin. The A domain PchD catalyzes the activation of salicylic acid (Sal) and its transfer onto the ArCP domain of PchE. (E) Biosynthetic pathway of the antitumor antibiotic vicenistatin. The A domain VinN catalyzes the activation of (2*S*,3*S*)-3- methylaspartic acid (2*S*,3*S*-MeAsp) and its transfer onto the CP VinL. The biosynthetic pathways and domains described in this study are shown in green, blue, orange, and yellow. Cy and E refer to the cyclization and epimerase domains, respectively.

Protein–protein interactions (PPIs) are key determinants of enzyme selectivity and pathway organization in NRPS systems.^4^ Structural studies of A–CP complexes have revealed specific PPIs that govern CP binding and substrate loading, allowing A domains to distinguish cognate from non-cognate CPs (Fig. 1A).^5^ Notably, A–CP interface features vary substantially among different pairs (Extended Data Fig. 1).^6–11^ Structural and mutational studies improved the interactions between BasE from *Acinetobacter baumannii* acinetobactin biosynthesis and EntB from *Escherichia coli* enterobactin biosynthesis.^6^ More recently, computational docking and mutagenesis enabled the engineering of a functional non-native PPI between PltF from *Pseudomonas fluorescens* pyoluteorin biosynthesis and the CP from *E. coli* fatty acid biosynthesis.^12^ These studies highlight the importance of appropriate recognition interfaces for high turnover in biosynthetic systems.^6,12^

Because A domains control substrate incorporation, they have been major targets in NRPS engineering, including A-domain substitution,^13^ A–CP didomain substitution,^14^ complete module exchange (C–A–CP tridomains),^15^ CP–C–A tridomain substitution,^16^ A-domain subdomain swapping,^17^ and active-site engineering.^18–21^ Despite these efforts,^22^ most combinatorial biosynthetic attempts have produced only small amounts of product or no detectable product.^23^ Recent *de novo* NRPS design based on exchange units and exchange-unit condensation (C) domains provides a notable exception.^24,25^ However, substitution-based systems may often lose the productive PPIs present in wild-type (wt) NRPSs. We therefore sought to establish a PPI- engineering strategy for designing customized NRPS systems, reasoning that combining PPI engineering with substitution-based approaches could enable the rational construction of customized NRPSs and the production of new peptides and peptide derivatives.

To test this concept, we selected enterobactin, vibriobactin, pyochelin, and vicenistatin biosynthetic pathways as model systems (Fig. 1B–E). Enterobactin is an *E. coli* siderophore biosynthesized by EntE, EntB, and EntF (Fig. 1B and Extended Data Fig. 2).^26^ EntE activates 2,3- dihydroxybenzoic acid (DHB) and transfers it to the Ppant thiol of the aryl CP (ArCP) domain of EntB, after which EntF catalyzes condensation with L-Ser, iterative elongation, and trilactone formation. Vibriobactin, produced by *Vibrio cholerae*, is assembled from three DHB molecules, two L-Thr molecules, and one norspermidine (NSPD) molecule. This assembly is carried out by VibE, VibB, VibF, and VibH through steps related to enterobactin biosynthesis (Fig. 1C).^27^ The reconstruction of VibE, VibB, and VibH generates a DHB–NSPD conjugate, and VibH accepts DHB-loaded EntB, which indicates native compatibility between EntB(ArCP) and VibH.^27,28^ Pyochelin, a salicylate (Sal)-capped siderophore from *Pseudomonas aeruginosa*, is derived from Sal and two L-Cys. Sal is activated by PchD and transferred to the *N*-terminal ArCP domain of PchE (Fig. 1D).^29^ Vicenistatin is a macrolactam antibiotic produced by *Streptomyces halstedii* (Fig. 1E).^30^ In its biosynthesis, VinN loads (2*S*,3*S*)-3-methylaspartic acid onto the CP VinL. After VinO-catalyzed decarboxylation, VinM loads L-Ala onto 3-aminoisobutyryl-VinL to generate dipeptidyl-VinL. Biosynthesis then proceeds through VinK, VinP1–P4, VinJ and VinC (Fig. 1E).^31^

## Results

### Structure-guided design of protein interfaces between adenylation and carrier protein domains

As an initial test case for engineering a productive non-cognate A–CP interaction, we selected the DHB-selective A domain EntE from enterobactin biosynthesis and the CP VinL from vicenistatin biosynthesis, with the aim of enabling transfer of DHB onto VinL (Fig. 1B,E,2A). To place VinL in the context of known ArCPs, we constructed a phylogenetic tree with VinL and representative ArCP domains (Extended Data Fig. 3). VinL shares 27% sequence identity with EntB(ArCP), and structural alignment showed an RMSD of 2.6 Å (Extended Data Fig. 3). To guide interface design, we compared the cognate EntE–EntB(ArCP) complex (PDB ID: 3RG2) and the VinM–VinL complex (PDB ID: 8K4R). Both have been structurally characterized and share a common mode of A–CP recognition (Extended Data Fig. 4).^6,11^ In both complexes, helix II of the CP engages the *N*-terminal A_core_ region of the partner A domain (Extended Data Fig. 4).^6,11^ Modeling of the non- cognate EntE–VinL interface suggested electrostatic incompatibility at this recognition surface (Fig. 2B). In the native EntE–EntB(ArCP) complex, Arg254 of EntB(ArCP) forms salt bridges with Glu292 of EntE (Fig. 2B). In VinL, the corresponding position is occupied by Glu45, suggesting an electrostatic mismatch with the native EntE surface. We therefore designed EntE E292K and E292R to increase the positive character of the EntE interface and promote productive VinL recognition. EntE E292K and E292R were prepared for experimental testing (Supplementary Fig. S1,S2). Recombinant wtEntE, VinL, and EntB(ArCP) were prepared according to previously reported methods,^19,32,33^ and EntB(ArCP) and VinL were converted into their *holo* forms using Sfp.^32,33^

**Fig. 2.**
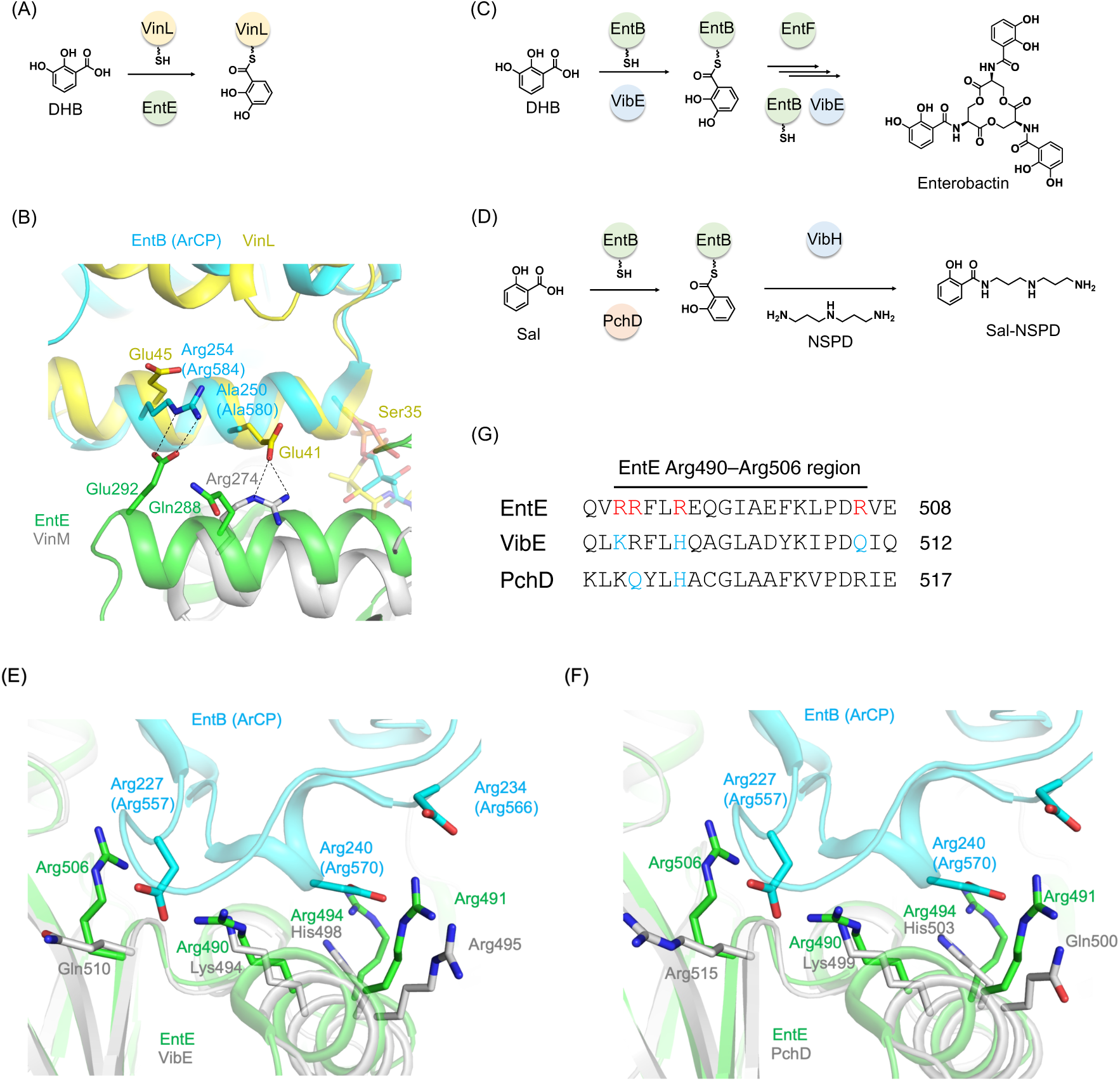
Structure-guided design of non-cognate adenylation–carrier protein interfaces for engineered NRPS pathways. (A) Design of a non-cognate EntE–VinL pair to enable the transfer of DHB from the adenylation domain EntE to the carrier protein VinL. (B) Structural model of the non-cognate EntE–VinL interface based on the cognate EntE–EntB(ArCP) and VinM–VinL complexes, highlighting residues targeted for reprogramming toward VinL recognition. Salt bridge interactions in the interfaces are shown as black dashed lines. EntE, EntB(ArCP), VinM and VinL are colored green, cyan, light gray, and yellow, respectively. Residue numbers in parentheses correspond to those of the EntE–EntB(ArCP) fusion protein, which was used for structural determination of the EntE–EntB(ArCP) complex (PDB ID: 3RG2). (C) Design of a chimeric enterobactin biosynthetic pathway by replacing wtEntE with VibE. (D) Design of a non- native biosynthetic pathway for the salicylic acid–norspermidine (Sal–NSPD) conjugate using PchD, EntB, and VibH. (E) Structural model of the non-cognate VibE–EntB(ArCP) interface based on the cognate EntE–EntB(ArCP) complex, highlighting residues targeted for reprogramming toward EntB(ArCP) recognition. The VibE structural model was generated by AlphaFold3. EntE, EntB(ArCP), and VibE are colored green, cyan, and light gray, respectively. (F) Structural model of the non-cognate PchD–EntB(ArCP) interface based on the cognate EntE– EntB(ArCP) complex, highlighting residues targeted for reprogramming toward EntB(ArCP) recognition. The PchD structural model was generated by AlphaFold3. EntE, EntB(ArCP), and PchD are colored green, cyan, and light gray, respectively. (G) Sequence alignment of the EntE Arg490–Arg506 region with the corresponding regions of VibE and PchD. Residues selected for interface reprogramming are highlighted.

We next investigated whether interface reprogramming could be applied to construct functional chimeric NRPS pathways. For this purpose, we selected the DHB-selective A domain VibE from vibriobactin biosynthesis, the Sal-selective A domain PchD from pyochelin biosynthesis, and EntB(ArCP) from enterobactin biosynthesis (Fig. 1B–D and Extended Data Fig. 2). Our hypothesis was that reprogramming the VibE–EntB(ArCP) and PchD–EntB(ArCP) interfaces would enable customized pathways: VibE–EntB–EntF for enterobactin biosynthesis and PchD– EntB–VibH for the production of a non-native Sal–NSPD conjugate (Fig. 2C,D). The native EntE–EntB(ArCP) structure reveals that the *C*-terminal A_sub_ region of EntE interacts with loop 1 of EntB (Leu226–Asp244) (Fig. 2E,F).^6^ Within this interface, EntE Arg490, Arg491, Arg494, and Arg506 form salt bridges with EntB(ArCP) Asp227, Asp234, and Asp240 (Fig. 2E,F).^6^ Consequently, we used this cognate interface as a template for reprogramming non-cognate A domains to recognize EntB(ArCP). Our focus was on the Lys494–Gln510 region of VibE and the Lys499–Gln515 region of PchD, which correspond to the EntE Arg490–Arg506 interface segment (Fig. 2E,F,G). To minimize perturbation of downstream pathway function, we conservatively introduced a limited number of EntE-like Arg residues. In VibE, Lys494, His498 and Gln510 were replaced with Arg residues corresponding to EntE Arg490, Arg494 and Arg506, respectively, generating H498R and Q510R single mutants, the H498R/Q510R double mutant and the K494R/H498R/Q510R triple mutant. In PchD, Gln500 and His503 were replaced with Arg residues corresponding to EntE Arg491 and Arg494, respectively, generating Q500R and H503R single mutants and the Q500R/H503R double mutant. All engineered VibE and PchD variants were prepared as *C*-terminally His_6_-tagged proteins (Supplementary Fig. S3–S9). Recombinant wtVibE, VibH, and wtPchD were prepared as previously reported.^27,29^

### Assessing the efficacy of interface reprogramming

We first evaluated DHB transfer by wtEntE and the PPI-reprogrammed EntE mutants E292K and E292R to the cognate CP EntB(ArCP) and the non-cognate CP VinL using time-course MALDI- TOF MS analysis (Fig. 3 and Supplementary Fig. S15–S20). wtEntE rapidly loaded EntB(ArCP), reaching near-complete conversion within 30 min, and both EntE E292K and E292R retained efficient loading of EntB(ArCP) (Fig. 3A–3C). By contrast, wtEntE generated only a small amount of DHB-loaded VinL even after 120 min, whereas both E292K and E292R converted *holo*-VinL almost completely to the DHB-loaded form within 120 min (Fig. 3D–F). Thus, charge reversal at the EntE interface enhanced productive DHB transfer to VinL while retaining activity toward the cognate partner EntB(ArCP).

**Fig. 3.**
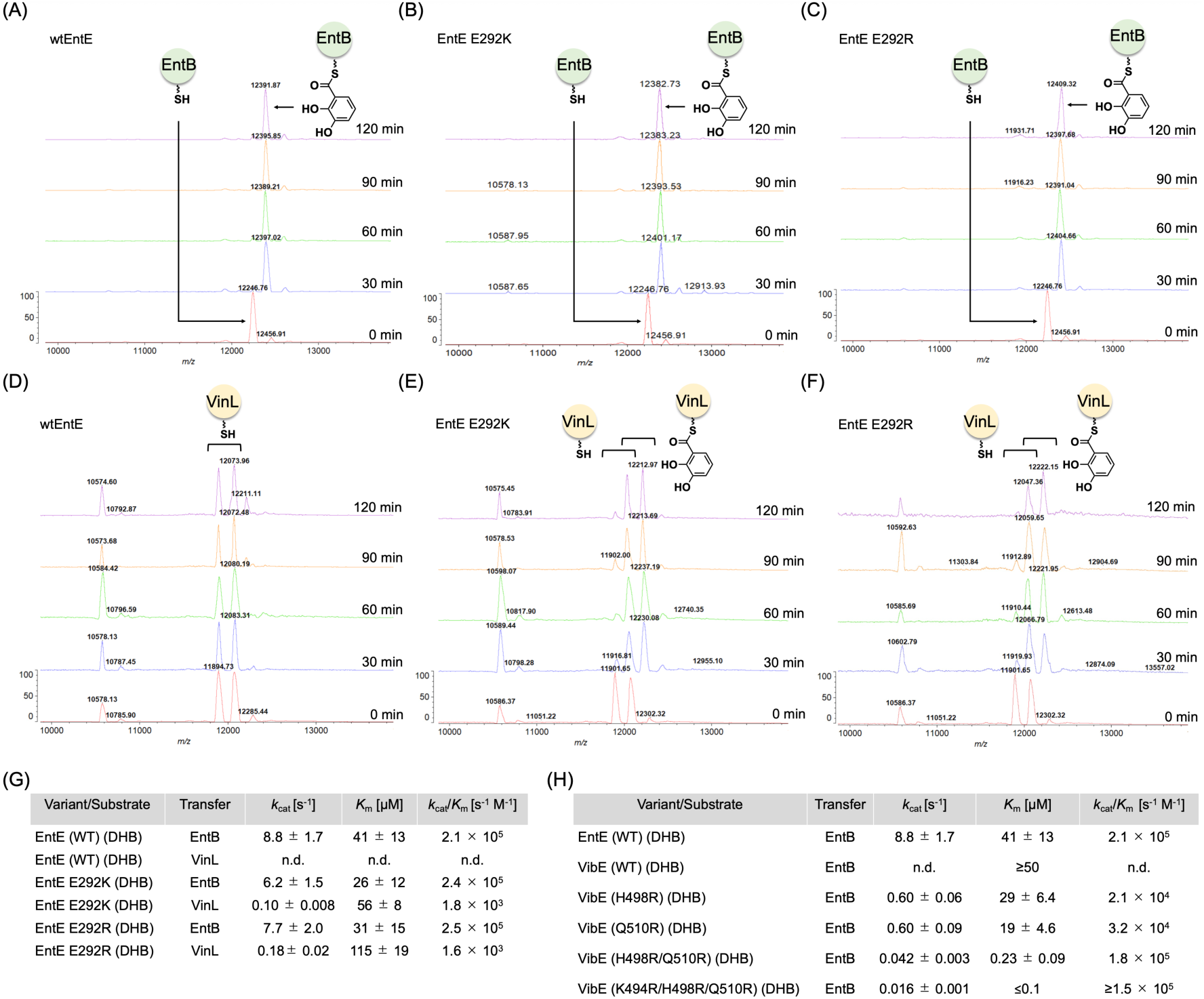
Interface reprogramming promotes DHB transfer to non-cognate carrier proteins (CPs). (A–C) Time-course MALDI-TOF mass spectra for DHB loading onto EntB(ArCP) by wtEntE (A), EntE E292K (B), and EntE E292R (C). Spectra were acquired at 0, 30, 60, 90, and 120 min. Peaks corresponding to the *holo* and DHB-loaded forms of EntB(ArCP) are indicated. (D–F) Time-course MALDI-TOF mass spectra for DHB loading onto VinL by wtEntE (D), EntE E292K (E), and EntE E292R (F). Spectra were acquired at 0, 30, 60, 90, and 120 min. Peaks corresponding to the *holo* and DHB-loaded forms of VinL are indicated. The +178-Da mass is likely derived from the spontaneous dephosphorylation of an α-*N*-phosphogluconoylation modification occurring on the His-tag of VinL in *E. coli*.^34^ (G) Steady-state kinetic parameters for DHB transfer by wtEntE and EntE interface mutants with the cognate CP EntB(ArCP) and the non-cognate CP VinL. No detectable activity was observed for wtEntE with VinL under the tested conditions. Data represent the mean of three separate measurements; error bars denote standard deviation. Apparent kinetic constants were determined by HPLC. (H) Steady-state kinetic parameters for DHB transfer by wtVibE and VibE interface mutants to the non-cognate CP EntB(ArCP). *k*_cat_, *K*_m_, and *k*_cat_/*K*_m_ could not be reliably estimated because wtVibE activity did not reach saturation over the tested EntB(ArCP) concentrations (0.2–50 μM). Accordingly, the apparent *K*_m_ for the ArCP substrate was judged to be ≥50 μM. For the K494R/H498R/Q510R triple mutant, the apparent *K*_m_ could not be accurately determined because of its very low value and was therefore estimated to be ≤ 0.1 μM. Apparent kinetic constants were determined by a malachite green colorimetric assay.^35^ Data represent the mean of three separate measurements, and error bars denote standard deviation. n.d.: not determinable. Expanded views of the MALDI- TOF mass spectra in Fig. 3A–F are provided in Supplementary Fig. S15–S20.

Next, we determined the steady-state kinetic parameters for DHB transfer to EntB(ArCP) and VinL (Fig. 3G and Extended Data Fig. 5). For EntB(ArCP), wtEntE, EntE E292K, and EntE E292R exhibited similar catalytic efficiencies, with *k*_cat_/*K*_m_ values of 2.1 × 10^5^, 2.4 × 10^5^, and 2.5 × 10^5^ s^-1^ M^-1^, respectively (Fig. 3G). No detectable DHB transfer from wtEntE to VinL was observed under these conditions. However, EntE E292K and E292R supported measurable transfer to VinL, with *k*_cat_/*K*_m_ values of 1.8 × 10^3^ and 1.6 × 10^3^ s^-1^ M^-1^, respectively (Fig. 3G). These data quantitatively confirm that interface reprogramming at position 292 enables productive DHB transfer to VinL while preserving efficient transfer to EntB(ArCP).

We then analyzed DHB transfer from wtVibE and interface-engineered VibE mutants to EntB(ArCP) (Fig. 3H and Extended Data Fig. 6). wtVibE showed no detectable transfer under these conditions, whereas the H498R and Q510R single mutants supported measurable activity, with *k*_cat_/*K*_m_ values of 2.1 × 10^4^ and 3.2 × 10^4^ s^-1^ M^-1^, respectively (Fig. 3H). The H498R/Q510R double mutant exhibited higher catalytic efficiency (*k*_cat_/*K*_m_ = 1.8 × 10^5^ s^-1^ M^-1^), primarily due to a substantially reduced apparent *K*_m_. The K494R/H498R/Q510R triple mutant showed comparable activity, with a *k*_cat_ of 0.016 ± 0.001 s^-1^ and an apparent *K*_m_ estimated to be below 0.1 µM (Fig. 3H). Across the mutant series, the apparent *K*_m_ decreased while *k*_cat_ tended to decrease, suggesting a trade-off between improved interprotein recognition and maximal catalytic turnover. Together, these results demonstrate that the progressive introduction of EntE-like interface residues substantially enhances productive DHB transfer from VibE to EntB(ArCP).

### NRPS pathway construction by interface reprogramming

Having established that non-cognate PPIs can be engineered, we next investigated whether interface reprogramming could construct functional non-native biosynthetic pathways through domain swapping. We began by reconstituting enterobactin biosynthesis *in vitro* using wtEntE, EntB, and EntF (Fig. 1B and Extended Data Fig. 2). Recombinant EntF was overproduced and converted to its *holo* form with Sfp, as described previously.^26^ Incubation with DHB, L-Ser, and ATP yielded the DHB–Ser monomer, dimer, and trimer, with *m/z* 242, 465, and 688 [(M + H)^+^], respectively, along with enterobactin with *m/z* 670 [(M + H)^+^] (Fig. 4A,C and Extended Data Fig. 2).

**Fig. 4.**
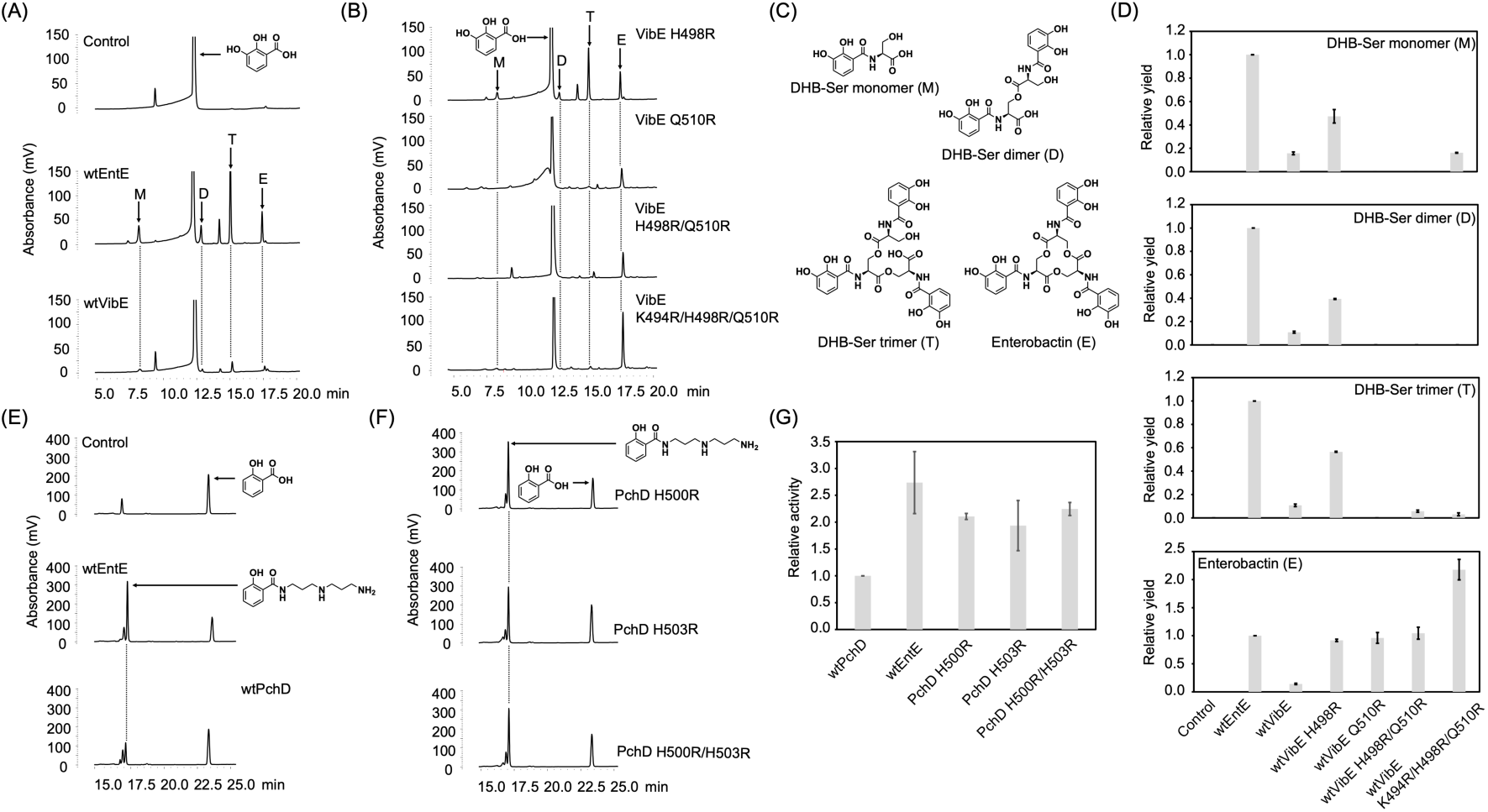
Reprogramming adenylation–carrier protein interfaces enables biosynthesis of native and non-native products. (A) HPLC chromatograms of *in vitro* reactions for enterobactin biosynthesis using wtEntE–EntB–EntF and wtVibE–EntB–EntF systems. Shown are a control reaction containing EntB and EntF with 2,3-dihydroxybenzoic acid (DHB) and L-Ser (top), a reaction containing wtEntE, EntB, and EntF with DHB and L-Ser (middle), and a reaction containing wtVibE, EntB and EntF with DHB and L-Ser (bottom). (B) HPLC chromatograms of *in vitro* reactions for enterobactin biosynthesis using VibE mutant–EntB–EntF systems. Reaction mixtures contained VibE H498R, VibE Q510R, VibE H498R/Q510R, and VibE K494R/H498R/Q510R, shown from top to bottom. (C) Structures of enterobactin and its early- stage intermediates. (D) Relative production of enterobactin and its early-stage intermediates by wtEntE–EntB–EntF, wtVibE–EntB–EntF, and VibE mutant–EntB–EntF systems. Values were normalized to the corresponding product levels obtained with wtEntE–EntB–EntF, which was set to 1 (*n* = 3 measurements). Error bars denote standard deviation. (E) HPLC chromatograms of *in vitro* reactions for synthesis of the non-native Sal–norspermidine (Sal–NSPD) conjugate using wtEntE–EntB–VibH and wtPchD–EntB–VibH systems. Shown are a control reaction containing EntB and EntF with Sal and NSPD (top), a reaction containing wtEntE, EntB, and VibH with Sal and NSPD (middle), and a reaction containing wtPchD, EntB, and VibH with Sal and NSPD (bottom). (F) HPLC chromatograms of *in vitro* reactions for synthesis of the non-native Sal– NSPD conjugate using PchD mutant–EntB–VibH systems. Reaction mixtures contained PchD Q500R (top), PchD H503R (middle), and PchD Q500R/H503R (bottom). (G) Relative initial velocities of Sal–NSPD conjugate formation by wtEntE–EntB–VibH, wtPchD–EntB–VibH, and PchD mutant–EntB–VibH systems. Values were normalized to the initial velocity obtained with the wtPchD–EntB–VibH system, which was set to 1 (*n* = 3 measurements). Error bars denote standard deviation.

To assess the impact of A-domain substitution in a full NRPS pathway, we replaced wtEntE with wtVibE in the enterobactin pathway (Fig. 1B,C,2B). Although the DHB–Ser monomer, dimer, trimer, and enterobactin were all detectable by HPLC, their production levels decreased to 16%, 11%, 11%, and 11%, respectively, relative to the native wtEntE–EntB–EntF pathway. This is consistent with the intrinsically poor substrate-transfer activity of wtVibE toward EntB(ArCP) (Fig. 3H,4A,D). Therefore, we investigated whether PPI reprogramming could restore pathway productivity. We introduced the reprogrammed VibE mutants H498R, Q510R, H498R/Q510R, and K494R/H498R/Q510R into enterobactin biosynthesis *in vitro* (Fig. 4B). Relative to the native wtEntE–EntB–EntF pathway, the H498R and K494R/H498R/Q510R pathways produced the DHB–Ser monomer in 47% and 16% yields, respectively, while no monomer was detected for Q510R or H498R/Q510R (Fig. 4B,D). The DHB–Ser dimer was detected only with H498R, in 39% yield. The trimer was produced by H498R, H498R/Q510R, and K494R/H498R/Q510R in 57%, 5.7%, and 3.1% yields, respectively, with no detectable trimer formation for Q510R (Fig. 4B,D). Notably, enterobactin production by the H498R, Q510R, H498R/Q510R, and K494R/H498R/Q510R pathways reached 0.91-, 0.96-, 1.0-, and 2.2-fold that of the native wtEntE–EntB–EntF pathway, respectively (Fig. 4B,D). Thus, despite lower or undetectable accumulation of early DHB-containing intermediates in some cases, interface-reprogrammed VibE variants restored enterobactin production to levels comparable to, or exceeding, that of the native pathway. These results indicate that simple domain substitution with non-cognate partners compromises pathway turnover, whereas combining domain substitution with PPI reprogramming restores pathway function, as exemplified by VibE K494R/H498R/Q510R– EntB–EntF (Fig. 4A,B,D).

We next tested whether this strategy could support the biosynthesis of a non-native product. Replacing wtVibE, VibB, and DHB with wtPchD, EntB, and Sal, respectively, was expected to enable the formation of *N*^1^-(2-hydroxybenzoyl) norspermidine (Sal–NSPD) by the wtPchD– EntB–VibH (Fig. 1B–D,2D). HPLC analysis showed that Sal–NSPD was not detected in the control reaction but was formed in the presence of either wtEntE–EntB–VibH or wtPchD–EntB–VibH (Fig. 4E). The product peak was more prominent in the wtEntE–EntB–VibH reaction than in the wtPchD–EntB–VibH reaction and was structurally confirmed by MS/MS analysis.^36^ Consistent with this result, the initial velocity of Sal–NSPD formation by the wtEntE–EntB–VibH pathway was 2.7-fold that of wtPchD–EntB–VibH (Fig. 4G and Extended Data Fig. 7). Interface reprogramming of PchD improved the productivity of the non-native pathway. PchD Q500R, H503R, and Q500R/H503R all supported Sal–NSPD formation (Fig. 4F) and increased the initial velocity to 2.1-, 1.9- and 2.2-fold that of wtPchD–EntB–VibH, respectively (Fig. 4G and Extended Data Fig. 7). The Q500R/H503R double mutant showed marginally higher activity than the single mutants and approached the activity of wtEntE–EntB–VibH, indicating substantial recovery of productive communication between PchD and EntB.

### Structural analysis of a reprogrammed interface

To understand the molecular basis for the reprogrammed interfaces and altered specificity, we attempted to determine the crystal structure of the VibE mutant–EntB(ArCP) complex. Because the transient VibE mutant–EntB(ArCP) complex is difficult to crystallize, we employed a cross- linking strategy^10,11,37–43^ to generate a covalent VibE mutant–EntB(ArCP) complex. We introduced an N239C mutation into each VibE mutant and performed cross-linking reactions with EntB(ArCP) using bromoacetyl pantetheinamide **C2Br** (see Supplementary Note). We then evaluated the cross-linking efficiency of each VibE mutant (Extended Data Fig. 8). Based on these results, we performed a large-scale cross-linking reaction of VibE N239C/K494R/H498R/Q510R with EntB(ArCP) in the presence of **C2Br**, and successfully determined the crystal structure of the VibE N239C/K494R/H498R/Q510R–EntB(ArCP) complex at 2.38 Å resolution (Fig. 5A and Extended Data Fig. 9; Table S1).

**Fig. 5.**
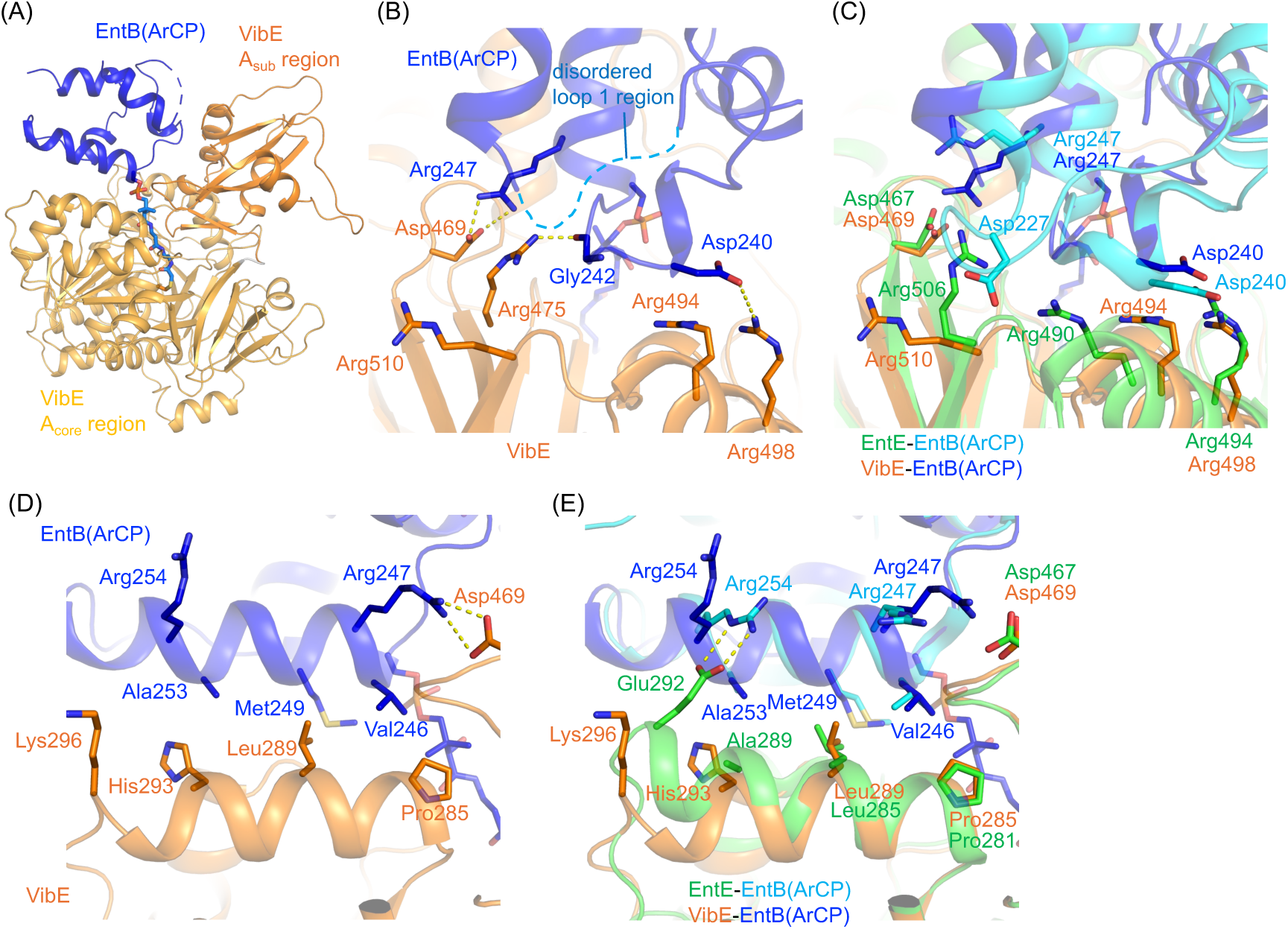
Structure of the VibE–EntB(ArCP) complex. (A) Overall structure of the VibE– EntB(ArCP) complex. The *N*-terminal A_core_ and C-terminal A_sub_ regions of VibE are colored bright orange and orange, respectively. EntB(ArCP) is colored blue. (B) Binding interface between the VibE A_sub_ region and EntB(ArCP) in the VibE–EntB(ArCP) complex. VibE and EntB(ArCP) are colored orange and blue, respectively. Salt bridges and a hydrogen bond are shown as yellow dashed lines. The disordered region (Asp227–Pro232) in loop 1 of EntB(ArCP) is shown as a cyan dashed line. (C) Superimposition of the A_sub_ region interfaces of the VibE–EntB(ArCP) complex and the EntE–EntB(ArCP) complex (PDB ID: 3RG2). VibE and EntB(ArCP) in the VibE–EntB(ArCP) complex are colored orange and blue, respectively. EntE and EntB(ArCP) in the EntE–EntB(ArCP) complex are colored green and cyan, respectively. (D) Binding interface between the VibE A_core_ region and EntB(ArCP) in the VibE–EntB(ArCP) complex. VibE and EntB(ArCP) are colored orange and blue, respectively. (E) Superimposition of the A_core_ region interfaces of the VibE–EntB(ArCP) complex and the EntE–EntB(ArCP) complex (PDB ID: 3RG2). VibE and EntB(ArCP) in the VibE–EntB(ArCP) complex are colored orange and blue, respectively. EntE and EntB(ArCP) in the EntE–EntB(ArCP) complex are colored green and cyan, respectively.

The interface area of the VibE–EntB(ArCP) complex is approximately 660 Å^2^, which is slightly smaller than that of the EntE–EntB(ArCP) complex (approximately 920 Å^2^). In the EntE– EntB(ArCP) complex structure, EntB(ArCP) interacts with both the *N*-terminal A_core_ (residues 1– 431) and *C*-terminal A_sub_ (residues 434–542) regions of EntE.^6^ In contrast, in the VibE– EntB(ArCP) complex structure, EntB(ArCP) interacts predominantly with the *C*-terminal A_sub_ (residues 434–542) region of VibE. The difference may partly result from the Arg substitutions introduced into the A_sub_ region. Although EntB(ArCP) interacts with a similar surface region of the A_sub_ region in both the VibE–EntB(ArCP) and EntE–EntB(ArCP) complexes, the detailed interaction patterns are not identical. Arg498 of VibE forms a salt bridge (3.2 Å) with Asp240 of EntB(ArCP), as observed for the interaction between Arg494 of EntE and Asp240 of EntB (ArCP) in the EntE–EntB(ArCP) complex structure (Fig. 5B). However, the side-chain orientations of Arg494 and Arg510 in VibE differ from those of the corresponding Arg490 and Arg506 residues in EntE, respectively (Fig. 5C). Arg490 and Arg506 of EntE interact with Asp227 of EntB(ArCP) in the EntE–EntB(ArCP) complex structure, whereas Asp227 of EntB(ArCP) was disordered in the VibE–EntB(ArCP) complex structure. The aliphatic portion of the VibE Arg494 side-chain, which corresponds to EntE Arg490, forms van der Waals contacts (3.3–4.0 Å) with Asp240 of EntB(ArCP). Additionally, the observed electron density for the Arg510 side-chain of VibE was weak, suggesting conformational flexibility of this side-chain, possibly due to the absence of stabilizing interactions with EntB(ArCP). Consequently, several interactions observed in the EntE–EntB(ArCP) complex structure were absent in the VibE–EntB(ArCP) complex structure. In addition to interactions involving the introduced Arg residues, other residues in the A_sub_ region of VibE also contribute to the interaction with EntB(ArCP). Asp469 of VibE forms salt bridges (2.7 Å and 3.4 Å) with Arg247 of EntB(ArCP) (Fig. 5B). Arg475 of VibE forms a hydrogen bond with the main-chain carbonyl oxygen of Gly242 of EntB(ArCP).

In the EntE–EntB(ArCP) complex structure, EntB(ArCP) also interacts with the *N*-terminal A_core_ region of EntE. This includes a salt bridge between Glu292 of EntE and Arg254 of EntB(ArCP) (Fig. 5E).^6^ However, VibE has Lys296 at the position corresponding to Glu292 of EntE, and the Arg254 side-chain of EntB(ArCP) is oriented away from VibE (Fig. 5D). This results in the absence of a corresponding salt bridge in the VibE–EntB(ArCP) complex structure. Instead, the aliphatic side-chains of Val246, Met249, and Ala253 of EntB(ArCP) form hydrophobic contacts (3.8 Å, 4.5 Å, and 3.6 Å) with Pro285, Leu289, and His293 of VibE, respectively. Most of these hydrophobic contacts are also observed in the EntE–EntB(ArCP) complex structure (Fig. 5E), indicating that the hydrophobic interaction pattern in the A_core_ region is largely conserved between the two complexes.

### Molecular dynamic (MD) simulations

To further investigate the interaction modes between VibE and EntB(ArCP), we performed triplicate 500-ns MD simulations using a complex model consisting of EntB(ArCP) carrying a functional phosphopantetheine arm and the VibE K494R/H498R/Q510R mutant, without a covalent linkage between the two proteins. During the MD simulations, EntB(ArCP) remained associated with the surface of VibE (Extended Data Fig. 10A). Asp469 and Arg498 of VibE formed stable salt bridges with Arg247 and Asp240 of EntB(ArCP), respectively (Fig. 6A–C), with high frequency (1.00 for Asp469–Arg247 and 0.74 for Arg498–Asp240; Extended Data Fig. 10B), which is consistent with the VibE–EntB(ArCP) complex structure (Fig. 5B).

**Fig. 6.**
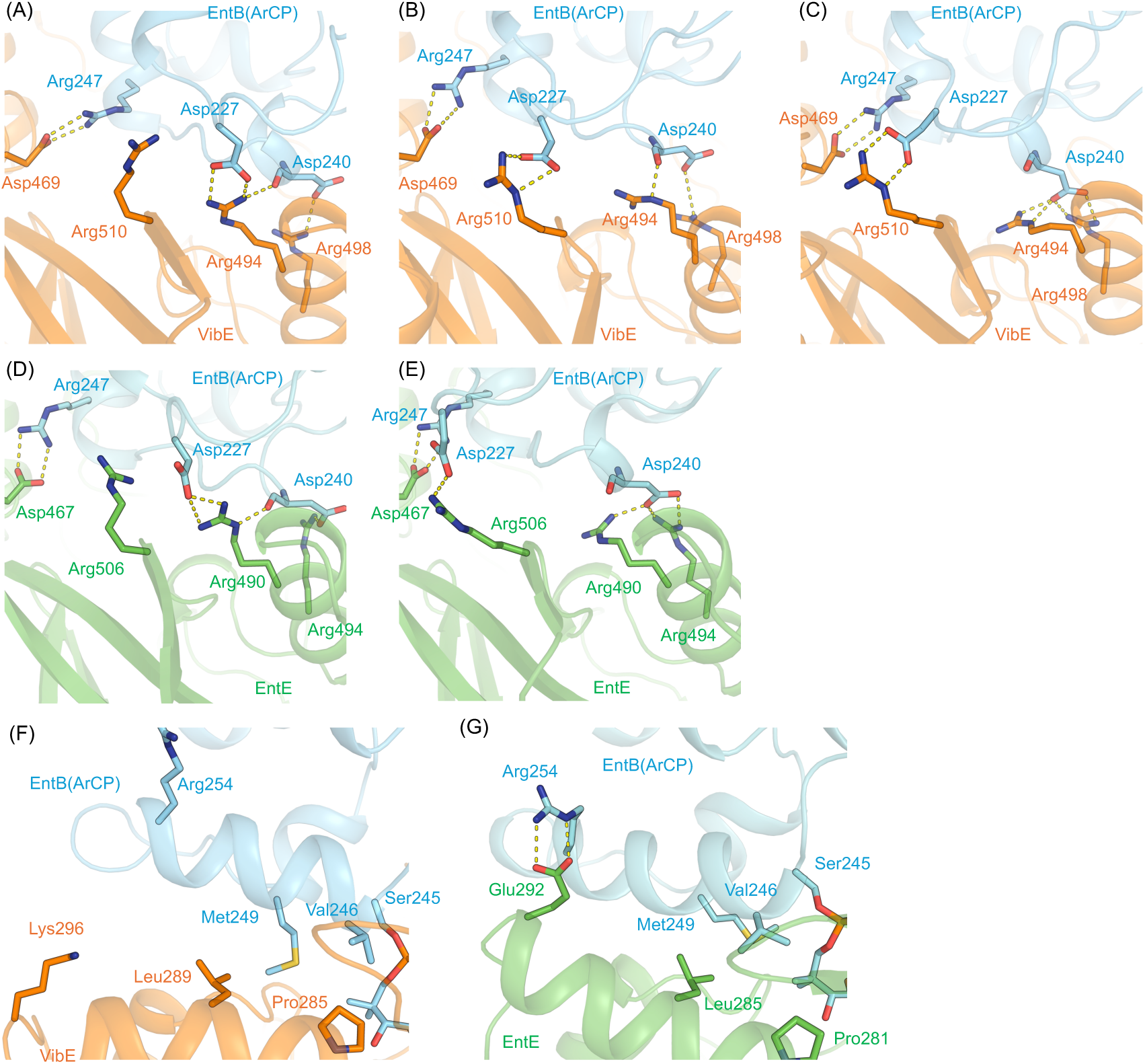
MD simulation analysis of the VibE–EntB(ArCP) and EntE–EntB(ArCP) complexes. (A–C) Binding interfaces between the VibE A_sub_ region and EntB(ArCP) in the VibE–EntB(ArCP) complex during MD simulation. Salt bridges and hydrogen bonds are shown as yellow dashed lines. Representative conformations are shown in which VibE Arg494 forms (A) salt bridges with EntB(ArCP) Asp227, (B) a hydrogen bond with the backbone amide of EntB(ArCP) Asp240, and (C) salt bridges with the side-chain of EntB(ArCP) Asp240. (D–E) Binding interfaces between the EntE A_sub_ region and EntB(ArCP) in the EntE–EntB(ArCP) complex during MD simulation. Representative conformations are shown in which EntE Arg490 forms (D) salt bridges with EntB(ArCP) Asp227 and a hydrogen bond with the backbone amide of EntB(ArCP) Asp240, and salt bridges with the side-chain of EntB(ArCP) Asp240. (F) Binding interface of the VibE A_core_ region and EntB(ArCP) in the VibE–EntB(ArCP) complex during MD simulation. (G) Binding interface of the EntE A_core_ region and EntB(ArCP) in the EntE–EntB(ArCP) complex during MD simulation.

Notably, the side-chain of Arg494 in VibE exhibited conformational flexibility during the simulations (Fig. 6A–C). Although Arg494 of VibE predominantly interacted with Asp240 of EntB(ArCP), as observed in the VibE–EntB(ArCP) complex structure, two major conformations were sampled: one in which the Arg494 side-chain formed a salt bridge with the Asp240 side-chain (Fig. 6C) and another in which it formed a hydrogen bond with the backbone carbonyl of Asp240 (Fig. 6B). Arg494 was also observed to reorient toward Asp227 of EntB(ArCP), forming a transient salt bridge (Fig. 6A), suggesting that multiple interaction modes are sampled in solution. The frequencies of these interactions for the Arg494 side-chain were as follows: 0.28 for the salt bridge with the Asp240 side-chain, 0.26 for the hydrogen bond with the Asp240 backbone, and 0.26 for the salt bridge with the Asp227 side-chain (Extended Data Fig. 10B). In addition, a salt bridge between VibE Arg510 and EntB(ArCP) Asp227 was frequently observed (frequency: 0.73; Fig. 6B–C and Extended Data Fig. 10B), despite its absence of this salt bridge in the VibE– EntB(ArCP) complex structure. The loop region containing Asp227 of EntB(ArCP) was disordered in the crystal structure, which may reflect crystal packing effects and/or the intrinsic flexibility of this loop region. Consistent with this interpretation, the loop 1 region containing Asp227, including both side-chain and backbone atoms, exhibited conformational variability during the MD simulations (Extended Data Fig. 10C,D).

To compare the interaction modes between VibE and EntB(ArCP) with those between EntE and EntB(ArCP), we performed triplicate 500-ns MD simulations using a model of the wtEntE– EntB(ArCP) complex. The salt bridges between EntE Asp467 and EntB(ArCP) Arg247, between EntE Arg494 and EntB(ArCP) Asp240, and between EntE Arg506 and EntB(ArCP) Asp227 were predominantly maintained during the MD simulations (frequencies: 1.00, 1.00, and 0.48; Fig. 6D–E and Extended Data Fig. 10B). The side-chain of EntE Arg490 exhibited conformational flexibility, similar to the corresponding VibE Arg494 residue. Although Arg490 of EntE predominantly interacted with Asp240 of EntB(ArCP) (Fig. 6E), this residue also transiently formed a salt bridge with Asp227 (Fig. 6D), consistent with the interaction observed in the EntE– EntB(ArCP) complex structure. Like VibE Arg494, EntE Arg490 sampled two major interaction modes with Asp240, involving either a side-chain salt bridge or a hydrogen bond to the Asp240 backbone carbonyl (Fig. 6D–E). The frequencies for these interactions of the Arg490 side-chain were 0.18 for the salt bridge with the Asp240 side-chain, 0.60 for the hydrogen bond with the Asp240 backbone, and 0.55 for the salt bridge with the Asp227 side-chain (Extended Data Fig. 10B). These results suggest that both EntE Arg490 and the corresponding VibE Arg494 dynamically switch between interactions with EntB(ArCP) Asp227 and Asp240 in solution.

The salt bridge between Glu292 of the EntE *N*-terminal A_core_ region and Arg254 of EntB(ArCP), observed in the EntE–EntB(ArCP) complex structure, remained stable during the MD simulations. In contrast, the A_core_ region of VibE did not form a salt bridge with Arg254 of EntB(ArCP) (Fig. 6F–G). Consistent with this, Arg254 of EntB(ArCP) remained oriented away from the VibE A_core_ region in representative conformations, never forming an EntE-like salt bridge (Extended Data Fig. 10E). Instead, Pro285 and Leu289 of VibE predominantly formed hydrophobic contacts with Val246 and Met249 of EntB(ArCP), respectively (Fig. 6F), similar to those observed in the simulations of the EntE–EntB(ArCP) complex. Overall, these MD simulations suggest that the EntE-like Arg substitutions in VibE reproduce the dynamic salt bridge network involving Asp227 and Asp240 of EntB(ArCP), resulting in an A_sub_ region interaction pattern that resembles that of EntE. However, the interaction pattern in the A_core_ region remains distinct because the EntE Glu292–Arg254 salt bridge interaction is not conserved in VibE.

## Discussion

Our results show that A–CP specificity can be rationally reprogrammed by minimal changes at the protein–protein interface. In the EntE–VinL system, charge reversal at a single interfacial position enabled robust DHB transfer to the non-cognate CP VinL while preserving efficient transfer to the cognate partner EntB(ArCP). This indicates that electrostatic complementarity is a major determinant of CP selectivity. Consistently, the progressive introduction of EntE-like residues into VibE reduced the apparent *K*_m_ for EntB(ArCP), supporting improved productive engagement of the CP. Although *k*_cat_ tended to decrease across the VibE mutant series, the overall gain in catalytic efficiency indicates that productive A–CP recognition is the dominant bottleneck in this non-cognate pair.

This principle was recapitulated at the pathway level. Replacing wtEntE with wtVibE strongly impaired enterobactin biosynthesis, consistent with the negligible DHB-transfer activity of wtVibE toward EntB(ArCP). In contrast, interface-engineered VibE variants restored pathway productivity to levels comparable to, or in one case exceeding, that of the native wtEntE–EntB– EntF pathway. Notably, this recovery occurred even when early DHB-containing intermediates were low or undetectable, suggesting that efficient pathway flux depends more on productive coupling between sequential catalytic steps than on the accumulation of upstream intermediates. The non-native Sal–NSPD pathway extended this conclusion beyond the restoration of a native product. Although PchD is the native Sal-activating enzyme, wtPchD–EntB–VibH was substantially less active than wtEntE–EntB–VibH, indicating that catalytic role alone does not guarantee productivity in a reconstituted non-native pathway. Reprogramming the PchD interface rescued this deficiency, with the Q500R/H503R double mutant approaching the activity of wtEntE–EntB–VibH, supporting the conclusion that the principal defect in the wtPchD–EntB pair is interfacial rather than catalytic.

Structural analysis provided detailed insights into engineered A–CP recognition. Although the crystal structure of the VibE–EntB(ArCP) complex did not fully reproduce the interaction pattern of the EntE–EntB(ArCP) complex, particularly the salt bridges involving EntB Asp227 (Fig. 5B,C), MD simulations revealed that the VibE–EntB(ArCP) interface samples multiple interaction modes in solution (Fig. 6). Some simulated conformations resembled the EntE– EntB(ArCP) interface, including transient salt bridges between VibE Arg494 or Arg510 and EntB Asp227. Thus, the crystal structure likely represents one accessible conformational state, potentially influenced by crystal packing and/or conformational stabilization during crystallization. The presence of multiple interaction modes in both VibE–EntB(ArCP) and EntE– EntB(ArCP) complexes suggests that productive A–CP recognition is mediated by a dynamic interaction network. Similar conformational heterogeneity has been reported for interactions between the ketosynthase-like decarboxylase (KS_Q_) domain and CP in the loading module of modular polyketide synthases,^44^ suggesting that dynamic interaction networks may be a common feature of CP-mediated PPIs.

These findings collectively identify A–CP interface compatibility as a dominant determinant of flux in engineered NRPS pathways. Minimal and targeted interface reprogramming can restore or create productive communication between non-cognate catalytic partners, providing a general framework for engineering new PPIs in assembly-line biosynthetic enzymes.

## Conclusion

Despite two decades of progress in NRPS research, domain substitution and module swapping still frequently yield inactive or poorly productive assembly lines because productive communication is lost between non-cognate catalytic partners. Here we show that this barrier can be overcome by structure-guided reprogramming of A–CP interfaces. We engineered EntE, VibE, and PchD to establish productive interactions with the non-cognate CPs VinL and EntB, generating the artificial PPIs EntE–VinL, VibE–EntB, and PchD–EntB, which were validated by biochemical, kinetic and structural analyses. Interface reprogramming enabled the construction of functional hybrid NRPS pathways spanning enterobactin, vicenistatin, vibriobactin, and pyochelin biosynthetic systems. Engineered VibE variants restored enterobactin production in non-native pathways to up to 2.2-fold that of the native EntE–EntB–EntF system, and engineered PchD variants enhanced production of a non-native Sal–NSPD conjugate with EntB and VibH. These results show that pathway output depends not only on catalytic competence but critically on compatibility at the A–CP interface. Structural analysis of the reprogrammed VibE–EntB complex, supported by MD simulations, showed that productive non-cognate recognition can be achieved by minimal rewiring of interfacial electrostatics while preserving the overall architecture of cognate A–CP assemblies. More broadly, our study identifies A–CP interface compatibility as a central determinant of NRPS engineering and provides a practical framework for constructing customized biosynthetic pathways to access new natural-product-inspired chemical space.

## Methods

### DNA construct design

The artificial genes encoding EntE (E292K, E292R, and N235C), VibE (H498R, Q510R, H498R/Q510R, K494R/H498R/Q510R, N239C, N239C/H498R, N239C/Q510R, N239C/H498R/Q510R, and N239C/K494R/H498R/Q510R), and PchD (Q500R, H503R, and Q500R/H503R) were synthesized by Eurofins Genomics (Tokyo, Japan). The synthesized *entE* genes were cloned into the expression vector pET28b using the NcoI and EcoRI restriction sites. The synthesized *vibE* genes were cloned into pET28b using the NcoI and SalI restriction sites, and the synthesized *pchD* genes were cloned into pET28b using the NcoI and XhoI restriction sites.

### Protein expression and purification

EntE and VibE variants were overproduced in *E. coli* BL21 (DE3) cells as *C*-terminally His_6_- tagged proteins. Overnight cultures were used to inoculate 1 L of Luria-Bertani (LB) medium supplemented with kanamycin (50 µg mL^-1^). Cultures were grown at 37 °C to an OD_600_ of 0.60– 0.80, induced with isopropyl β-D-1-thiogalactopyranoside (IPTG; final concentration of 0.1 mM), and incubated for an additional 3 h at 37 °C. Cells were harvested by centrifugation, resuspended in lysis buffer (20 mM Tris–HCl, pH 8.0, 0.5% Triton-X), and disrupted by sonication at 4 °C using an ultrasonic disruptor (UD201, Tomy Digital Biology Co., Ltd, Japan). After centrifugation, the supernatants were applied to a Ni-NTA agarose column (Qiagen, Germany), and bound proteins were eluted with an imidazole gradient. Eluted fractions were analyzed by SDS-PAGE with Coomassie Brilliant Blue staining, and protein concentrations were quantified using the Bradford assay.^45^ Fractions containing recombinant proteins were pooled, dialyzed against 20 mM Tris–HCl, pH 8.0, 1 mM MgCl_2_, and 1 mM TCEP, supplemented with glycerol to 10% (v/v), and stored at −80 °C.

Recombinant wtEntE and *holo*-ArCP were expressed and purified as previously reported.^26,33^ Recombinant *apo*- and *holo*-VinL were overproduced and purified as previously described.^32^ CoaA, CoaD, and CoaE were prepared as maltose-binding protein-fused recombinant proteins according to previously described methods.^46^ Sfp from *Bacillus subtilis* was prepared as a His-tagged recombinant protein as previously reported.^46^

### Transfer of DHB to the ArCP domain of EntB catalyzed by wtEntE and the mutants

Reaction mixtures (50 µL) contained recombinant *holo*-ArCP (10 µM), adenylation enzymes (500 nM wtEntE, 500 nM EntE E292R, or 500 nM EntE E292K), DHB (1 mM), and ATP (2.5 mM) in a buffer consisting of 75 mM Tris–HCl (pH 7.5), 10 mM MgCl_2_, and 5 mM DTT. In all experiments, the total DMSO concentration was kept at 1.0% (v/v). After the addition of all components, the reactions were incubated at 37 °C for 30, 60, 90, or 120 min, precipitated with acetone, resolubilized in MilliQ water, and subjected to matrix-assisted laser desorption/ionization time-of-flight mass spectrometry (MALDI-TOF-MS) analysis. All experiments were performed in duplicate.

### Transfer of DHB to VinL catalyzed by wtEntE and its mutants

Reaction mixtures (50 µL) contained recombinant *holo*-VinL (10 µM), adenylation enzymes (500 nM wtEntE, 500 nM EntE E292R, or 500 nM EntE E292K), DHB (1 mM), and ATP (2.5 mM) in a buffer consisting of 50 mM HEPES–NaOH (pH 8.0), 10 mM MgCl_2_, and 5 mM DTT. In all experiments, the total DMSO concentration was kept at 1.0% (v/v). After adding all components, the reactions were incubated at 37 °C for 30, 60, 90, or 120 min, precipitated with acetone, resolubilized in MilliQ water, and subjected to MALDI-TOF-MS analysis. All experiments were performed in duplicate.

### Determination of kinetic parameters of transfer reactions by HPLC

#### Transfer of DHB to the ArCP domain of EntB catalyzed by wtEntE and the mutants

Reaction mixtures (20 µL) contained recombinant *holo*-ArCP (1, 5, 10, 20, and 40 µM), adenylation enzymes (100 nM wtEntE, 100 nM EntE E292R, or 100 nM EntE E292K), DHB (1.25 mM), and ATP (3.1 mM) in a buffer consisting of 75 mM Tris–HCl (pH 8.0), 10 mM MgCl_2_, and 5 mM DTT. In all experiments, the total DMSO concentration was kept at 2.5% (v/v). Reaction mixtures were incubated at 37 ℃, and after 1 min the reactions were quenched by adding CH_3_CN (20 μL). The resulting mixtures were centrifuged at 20,000× *g* for 5 min at 4 °C, and aliquots (20 μL) were analyzed by HPLC. All experiments were performed in triplicate. Initial velocities were determined from the decrease in *holo*-ArCP, which was quantified using calibration curves generated with authentic *holo*-ArCP. Apparent kinetic constants were obtained by fitting the data to the Michaelis–Menten equation.

#### Transfer of DHB to VinL catalyzed by wtEntE and the mutants

Reaction mixtures (20 µL) contained recombinant *holo*-VinL (10, 20, 30, 40, 50, or 100 µM), adenylation enzymes (200 nM wtEntE, 200 nM EntE E292R, or 200 nM EntE E292K), 1.25 mM DHB, and 3.1 mM ATP. The buffer consisted of 50 mM HEPES–NaOH (pH 8.0), 10 mM MgCl_2_, and 5 mM DTT. In all experiments, the total DMSO concentration was maintained at 2.5% (v/v). Reaction mixtures were incubated at 37 ℃. After 30 min, reactions were quenched by adding 20 μL CH_3_CN. The resulting mixtures were centrifuged at 20,000× *g* for 5 min at 4 °C, and 20 μL aliquots were analyzed by HPLC. All experiments were performed in triplicate. Initial velocities were determined from the decrease in *holo*-VinL, quantified using calibration curves generated with authentic *holo*-VinL. Apparent kinetic constants were obtained by fitting the data to the Michaelis–Menten equation.

#### HPLC settings

Cosmosil Protein-R (Nacalai Tesque, Japan), φ4.6 × 250 mm; CH_3_CN/H_2_O containing 0.1% TFA (10% CH_3_CN at 0 min, 10–100% CH_3_CN over 30 min, 100% CH_3_CN over 40 min, 10% CH_3_CN over 50 min); 1 mL/min; 220 nm.

### Determination of kinetic parameters of transfer reactions by a malachite green phosphate assay

#### Transfer of DHB to the ArCP of EntB catalyzed by wtVibE and its mutants

Reaction mixtures (40 µL) contained recombinant *holo*-EntB(ArCP) (wtVibE: 10, 20, 30, 40, 50 μM; VibE H498R: 1, 5, 10, 20, 40, 50 μM; VibE Q510R: 1, 5, 10, 15, 20 μM; VibE H498R/Q510R: 0.2, 0.4, 0.8, 1, 5 μM; VibE K494R/H498R/Q510R: 0.01, 0.1, 0.5, 1 μM), adenylation enzymes (100 nM wtVibE, 100 nM VibE H498R, 100 nM VibE Q510R, 500 nM VibE H498R/Q510R, or 2 μM VibE K494R/H498R/Q510R), DHB (1 mM), ATP (200 µM), 150 mM hydroxylamine (pH 7.0), and 0.04 U inorganic pyrophosphatase (Sigma–Aldrich, USA, I1643) in a buffer consisting of 75 mM Tris–HCl (pH 7.5), 10 mM MgCl_2_, and 5 mM DTT. Working stocks of hydroxylamine were prepared fresh by combining 500 µL of 4 M hydroxylamine, 250 µL of water and 250 µL of 7 M NaOH on ice. In all experiments, the total DMSO concentration was kept at or below 1.25% (v/v). The reactions (40 µL) were run in 96-well half-area plates (Corning, USA, 3881). Reaction mixtures were incubated at 37 ℃, and after 10–20 min the reaction was quenched by adding 10 µL of working reagent from a malachite green phosphate assay kit (BioAssay Systems, USA). After 10 min incubation at room temperature, absorbance at 620 nm (A_620_) was measured on a MULTISKAN FC (Thermo Fisher Scientific). The A_620_ value of the reaction mixture in the absence of substrate was subtracted from the A_620_ value of a reaction mixture in the presence of substrate to estimate the adenylation activities. All experiments were performed in at least duplicate.

### *In vitro* enzymatic synthesis of enterobactin

Reaction mixtures (50 μL) contained 10 mM ATP, 1 mM DHB, 2 mM L-Ser, 100 nM adenylation enzymes (wtEntE, VibE H498R, VibE Q510R, VibE H498R/Q510R, and VibE K494R/H498R/Q510R), 10 μM *holo*-EntB, and 2 μM *holo*-EntF in 50 mM HEPES–NaOH (pH 8.0), 10 mM MgCl_2_, and 5 mM DTT. The mixtures were preincubated at 37 ℃ for 5 min, prior to the addition of L-Ser and *holo*-EntF, to allow acylation of *holo*-EntB with DHB by the adenylation enzymes. After L-Ser and *holo*-EntF were added, the reactions were incubated at 37 °C. After 30 min, 150 μL of 1 M HCl was added to quench the reactions. The acidified reaction mixtures were extracted with 1 mL of EtOAc, and the organic layers were evaporated to dryness. The resulting residue was dissolved in 50 μL of 50% (v/v) acetonitrile/water, and 20 μL aliquots were analyzed by HPLC. Control reactions, performed under identical conditions, omitted the adenylation enzymes. All experiments were performed in triplicate. HPLC settings: Cosmosil MS-II (Nacalai Tesque, Japan), φ4.6 × 250 mm; CH_3_CN /H_2_O containing 0.1% TFA (10% CH_3_CN at 0 min, 10–100% CH_3_CN over 30 min, 100% CH_3_CN over 40 min, 10% CH_3_CN over 50 min); 1 mL/min; 254 nm. LC-MS settings: Cosmosil MS-II (Nacalai Tesque, Japan), φ3.0 × 150 mm; CH_3_CN/H_2_O containing 0.1% formic acid (10% CH_3_CN at 0 min, 10–100% CH_3_CN over 30 min, 100% CH_3_CN over 40 min, 10% CH_3_CN over 50 min); 0.2 mL/min; 210 and 254 nm.

### *In vitro* enzymatic synthesis of *N*^1^-(2-hydroxybenzoyl)norspermidine

Reaction mixtures (50 μL) contained 5 mM ATP, 1 mM Sal, 5 mM NSPD, 1 μM adenylation enzymes (wtEntE, wtPchD, PchD Q500R, PchD H503R, and PchD Q500R/H503R), and 10 μM *holo*-EntB in 50 mM HEPES–NaOH (pH 8.0), 10 mM MgCl_2_, and 5 mM DTT. Prior to the addition of NSPD and 1 μM VibH, the reaction mixtures were preincubated at 37 °C for 10 min to allow acylation of *holo*-EntB with Sal by the adenylation enzymes. Following the addition of NSPD and VibH, the reactions were incubated at 37 °C. After 4 h, the reactions were quenched by the addition of CH_3_OH (50 μL). The resulting mixtures were centrifuged at 20,000×*g* for 5 min at 4 °C, and aliquots (50 μL) were analyzed by HPLC. Control reactions were performed under identical conditions, except that adenylation enzymes were omitted. All experiments were performed in duplicate.

HPLC settings: Cosmosil PAQ (Nacalai Tesque, Japan), φ4.6 × 250 mm; CH_3_CN /H_2_O containing 0.1% TFA (0% CH_3_CN 0–5 min, 0–100% CH_3_CN 5–30 min, 100% CH_3_CN 30–40 min, 0% CH_3_CN 40–50 min); 1 mL/min; 254 nm.

### Initial velocity analysis of *N*^1^-(2-hydroxybenzoyl)norspermidine formation

Reaction mixtures (50 μL) contained 5 mM ATP, 1 mM Sal, 5 mM NSPD, 1 μM adenylation enzymes (wtEntE, wtPchD, PchD Q500R, PchD H503R, and PchD Q500R/H503R), and 10 μM *holo*-EntB in 50 mM HEPES–NaOH (pH 8.0), 10 mM MgCl_2_, and 5 mM DTT. Before the addition of NSPD and 1 μM VibH, the reaction mixtures were preincubated at 37 °C for 10 min to allow acylation of *holo*-EntB with Sal by the adenylation enzymes. Following the addition of NSPD and VibH, the reactions were incubated at 37 °C. After 0, 30, 60, 90, and 120 min for PchD Q500R and PchD Q500R/H503R, and 0, 60, 120, 180, and 240 min for wtEntE, wtPchD, and PchD H503R, the reactions were quenched by adding CH_3_OH (50 μL). The resulting mixtures were centrifuged at 20,000×*g* for 5 min at 4 °C, and aliquots (50 μL) were analyzed by HPLC. All experiments were performed in triplicate.

HPLC settings: Cosmosil PAQ (Nacalai Tesque, Japan), φ4.6 × 250 mm; CH_3_CN/H_2_O containing 0.1% TFA (0% CH_3_CN 0–5 min, 0–100% CH_3_CN 5–30 min, 100% CH_3_CN 30–40 min, 0% CH_3_CN 40–50 min); 1 mL/min; 254 nm.

### Site-selective crosslinking reactions with a bromoacetyl pantetheinamide

#### Crosslinking of EntB(ArCP) and adenylation enzymes

Reaction mixtures (50 μL) contained 8 mM ATP, 15 mM MgCl_2_, 1 mM **C2Br**, 40 μM *apo*-EntB(ArCP), 0.01 μg/μL CoaA, 0.01 μg/μL CoaD, 0.01 μg/μL CoaE, and 0.01 μg/μL Sfp in 50 mM potassium phosphate (pH 7.0). Controls were treated with DMSO only (vehicle), and in all reactions the final DMSO concentration was maintained at 1% (v/v). Prior to the addition of adenylation enzymes (5 μM EntE N235C, VibE N239C, VibE N239C/H498R, VibE N239C/Q510R, VibE N239C/H498R/Q510R, or VibE N239C/K494R/H498R/Q510R), reaction mixtures were preincubated at 37 ℃ for 60 min to load **C2Br** onto EntB(ArCP). After the addition of adenylation enzymes, reactions were incubated at 37 ℃ for 16 h. Reactions were then treated with 5× SDS-loading buffer (strong reducing conditions) and analyzed by SDS-PAGE.

#### Crosslinking efficiency and validation of cysteine reactivity

Reaction mixtures (50 μL) contained 8 mM ATP, 15 mM MgCl_2_, 1 mM **C2Br**, 40 μM *apo*-EntB(ArCP), 0.01 μg/μL CoaA, 0.01 μg/μL CoaD, 0.01 μg/μL CoaE, and 0.01 μg/μL Sfp in 50 mM potassium phosphate (pH 7.0). Controls were treated with DMSO only (vehicle), and in all reactions, the final DMSO concentration was maintained at 1% (v/v). Prior to the addition of adenylation enzymes (5 μM wtEntE, EntE N235C, VibE K494R/H498R/Q510R, or VibE N239C/K494R/H498R/Q510R), reaction mixtures were preincubated at 37 ℃ for 60 min to load **C2Br** onto EntB(ArCP). After the addition of adenylation enzymes, reactions were incubated at 37 ℃ for 16 h. Reactions were treated with 5× SDS-loading buffer (strong reducing conditions) and subjected to SDS-PAGE.

#### Concentration dependence on EntB(ArCP)

Reaction mixtures (50 μL) contained 8 mM ATP, 15 mM MgCl_2_, 1 mM **C2Br**, 10−40 μM *apo*-EntB(ArCP), 0.01 μg/μL CoaA, 0.01 μg/μL CoaD, 0.01 μg/μL CoaE, and 0.01 μg/μL Sfp in 50 mM potassium phosphate (pH 7.0). In all reactions, the final DMSO concentration was maintained at 1% (v/v). Prior to the addition of 5 μM VibE N239C/K494R/H498R/Q510R, reaction mixtures were preincubated at 37 ℃ for 60 min to load **C2Br** onto EntB(ArCP). After the addition of the adenylation enzyme, reactions were incubated at 37 ℃ for 16 h. Reactions were treated with 5× SDS-loading buffer (strong reducing conditions) and subjected to SDS-PAGE.

#### Preparation and purification of the cross-linked VibE N239C/K494R/H498R/Q510R– EntB(ArCP) complex

For structural analysis of the VibE–EntB(ArCP) complex, pET26b-*vibE N239C/K494R/H498R/Q510R* was constructed. VibE fragments were amplified by PCR using pET28b-*vibE N239C/K494R/H498R/Q510R* as a template and the following primers: 5’- AAGGTCCTATGACAACCGATTTTACCCCTTGG -3’ (forward) and 5’- GGATCTCACTCGAGTGCGGCCGCAAGCTTG-3’ (reverse). The linearized vector DNA was amplified by PCR using the pET26b plasmid as a template and the following primers: 5’- CACTCGAGTGAGATCCGGCTGCTAACAAAG -3’ (forward) and 5’- GTTGTCATAGGACCTTGAAAAAGAACTTCAAGG -3’ (reverse). The amplified VibE fragments were inserted into the linearized vector DNA using the In-Fusion HD Cloning Kit (Clontech Laboratories, USA) and transformed into *E. coli* DH5α. The expression plasmid was verified by DNA sequencing (Azenta Life Sciences, USA).

Next, *E. coli* BL21(DE3) cells harboring pET26b-*vibE N239C/K494R/H498R/Q510R* were grown at 37 °C in LB broth containing 100 mg/L kanamycin. When the optical density at 600 nm reached 0.5, protein expression was induced by adding IPTG to a final concentration of 0.1 mM, and the cells were cultured further at 15 °C for 20 h. Recombinant VibE N239C/K494R/H498R/Q510R mutant protein, collected from cell-free extracts prepared by sonication, was purified using a Ni cOmplete His-tag affinity column (Sigma–Aldrich). The N-terminal His-tag of the VibE N239C/K494R/H498R/Q510R mutant was subsequently removed by treatment with HRV 3c protease to obtain the His-tag free VibE N239C/K494R/H498R/Q510R mutant. The recombinant EntB(ArCP) protein, comprising the Ile211–Lys285 region of EntB and fused to an N-terminal His-tag, was also prepared as described previously.^47^

For the large-scale cross-linking reaction, 200 µM His-tagged EntB(ArCP) was mixed with 600 µM pantetheineamide probe **C2Br**, 15 mM MgCl_2_, 10 mM ATP, 1.5 µM CoaA, 2.1 µM CoaD, 1.8 µM CoaE, and 7.5 µM Sfp in a buffer containing 50 mM HEPES-Na (pH 7.0) and 10% (v/v) glycerol, to a total solution volume of 0.96 mL. After incubation at 28 °C for 3 h, the cross-linking reaction was initiated by directly adding 50 µM His-tag free VibE N239C/K494R/H498R/Q510R mutant to the reaction mixture. The total solution volume was increased to 1.92 mL, and the EntB(ArCP) concentration was diluted to 100 µM by the addition of the VibE mutant solution. After incubation at 28 °C for 20 h, the generated VibE–EntB(ArCP) complex was purified by Ni affinity chromatography. The VibE–EntB(ArCP) complex was further purified using a Superdex 200 Increase 10/300 column (Cytiva, USA) equilibrated with 50 mM HEPES-Na (pH 8.0) and 100 mM NaCl. Finally, the VibE–EntB(ArCP) complex was desalted and concentrated using an Amicon Ultra centrifugal filter (Millipore, USA).

### Crystallization, data collection, and structural determination

Crystals of the VibE–EntB(ArCP) complex were grown at 4 °C using the sitting-drop vapor diffusion method. Protein solution [3 mg/mL in 10 mM Tris-HCl (pH 8.0), 100 mM NaCl] was mixed with an equal volume of reservoir solution [100 mM Tris-HCl (pH 8.5), 200 mM MgCl_2_ and 20% (v/v) PEG8000]. Crystals were flash-frozen in liquid nitrogen before X-ray data collection. Diffraction data were collected at beamline BL45XU at SPring-8 (Hyogo, Japan) and then processed with XDS software.^48^ The initial phase was determined using the Phaser program^49^ with the VibE–EntB(ArCP) complex model structure predicted by AlphaFold3^50^, as the search model. Coot was used for manual rebuilding of the model.^51^ Refmac was used to refine the structure.^52^ Molecular graphic images were prepared using PyMOL (https://pymol.org/2/). The geometries of the final VibE–EntB(ArCP) complex structure were evaluated using the program MolProbity^53^. The atomic coordinates and structure factors have been deposited in the Protein Data Bank (PDB entry XXXX).

### MD simulations

MD simulations were performed for the wtEntE–EntB(ArCP) and VibE K494R/H498R/Q510R– EntB(ArCP) complexes. Starting models were prepared from corresponding experimental structures, with disordered regions modeled using AlphaFold3^50^ predictions where necessary. EntB(ArCP) was modeled with a phosphopantetheinylated Ser245. The N239C crosslinking mutation in the VibE–EntB(ArCP) structure was reverted to Asn, and ADP was removed. Protonation states were assigned for pH 7.5 using Reduce^54^ and PROPKA3,^55^ followed by the addition of hydrogen atoms. Systems were prepared with the AMBER ff19SB^56^ protein force field, GAFF2 atom types for non-standard groups^57^, and the OPC water model.^58^ The phosphopantetheinylated Ser245 residue was described using RESP charges and parameters for phosphopantetheinyl-serine derived from a previously reported pantetheine force-field library.^59^ Hydrogen addition, solvation, ion placement, and Amber topology generation were performed with tleap in AmberTools 25.^60^ Each complex was solvated in an approximately 95 × 95 × 95 Å cubic box. KCl ion numbers were determined using SLTCAP,^61^ and each system was neutralized with counterions. After energy minimization and staged positional-restraint equilibration for 10 ns, triplicate 500 ns production simulations were performed for each complex at 310.15 K using GROMACS 2025.1.^62^ Trajectories were processed for periodic-boundary-condition correction before analysis. Backbone RMSD and Cα RMSF were calculated with gmx rms and gmx rmsf, respectively. Residue-pair heavy-atom contacts were defined using a minimum-distance cutoff of ≤ 0.45 nm, and salt bridges were defined using a minimum distance of ≤ 0.40 nm between standard charged atoms of oppositely charged residues. Hydrogen bonds were calculated with gmx hbond using the default geometric criteria.

### Data availability

The data supporting the findings of this study are available within the article and its Supplementary Information. The electron density map and atomic coordinates for the VibE– EntB(ArCP) complex crystal structure have been deposited in the PDB under accession code XXX.

## Supporting information

Supplementary Notes, and Supplementary Figure 1-21, Supplementary Table 1, and Supplementary References

## Acknowledgements

This work was supported by Adaptable and Seamless Technology transfer Program through Target-driven R&D (A-STEP) from Japan Science and Technology Agency (JST) Japan (Grant Number JPMJTR25U3 to F.I.). Additional support was provided by the Asahi Glass Foundation to F.I., the Institute for Fermentation, Osaka (IFO) to F.I., the 2025 Kindai University Research Enhancement Grants (IP002) to F.I., the Anti-aging Project for Private Universities to G.T., and the Japan Society for the Promotion of Science (25K01946 to A.M. and 22H05126 to T.T.). Further support came from the Research Support Project for Life Science and Drug Discovery (Basis for Supporting Innovative Drug Discovery and Life Science Research (BINDS)) from AMED under Grant Number JP25ama121027 to T.T., and JST SPRING under Grant Number JPMJSP2108 to T.U. We thank the staff of SPring-8 for assistance with X-ray data collection. Plasmids encoding EntE, CoaA, CoaD, CoaE, and Sfp were provided by Prof. Michael D. Burkart (University of California, San Diego).

## Author contributions

F.I., G.T., and A.M. conceived and supervised the project. F.I. and A.M. designed the experiments. F.I., K.K., and Y.N. performed protein preparation, biochemical assays, kinetic analyses, *in vitro* pathway reconstitution and crosslinking experiments. F.I. and A.M. designed the interface mutations and interpreted the biochemical data. I.A. and K.S. prepared crosslinked complexes for structural studies. I.A., K.S., and A.M. determined and analyzed the crystal structure. T.U. and T.T. performed molecular dynamics simulations, and T.U. and T.T. analyzed the simulation data. F.I. synthesized a chemical probe. F.I., T.U., T.K., F.K., T.E., T.T., S.F., G.T., and A.M. wrote the manuscript.

## Additional information

Extended data is available for this paper at XXX.

**Supplementary information** The online version contains supplementary material available at XXX.

**Correspondence and requests for materials** should be addressed to Fumihiro Ishikawa, Genzoh Tanabe, or Akimasa Miyanaga.

## Peer review information **XXX.**

Reprints and permission information is available at XXX.

---

**Extended Data Fig. 1.**
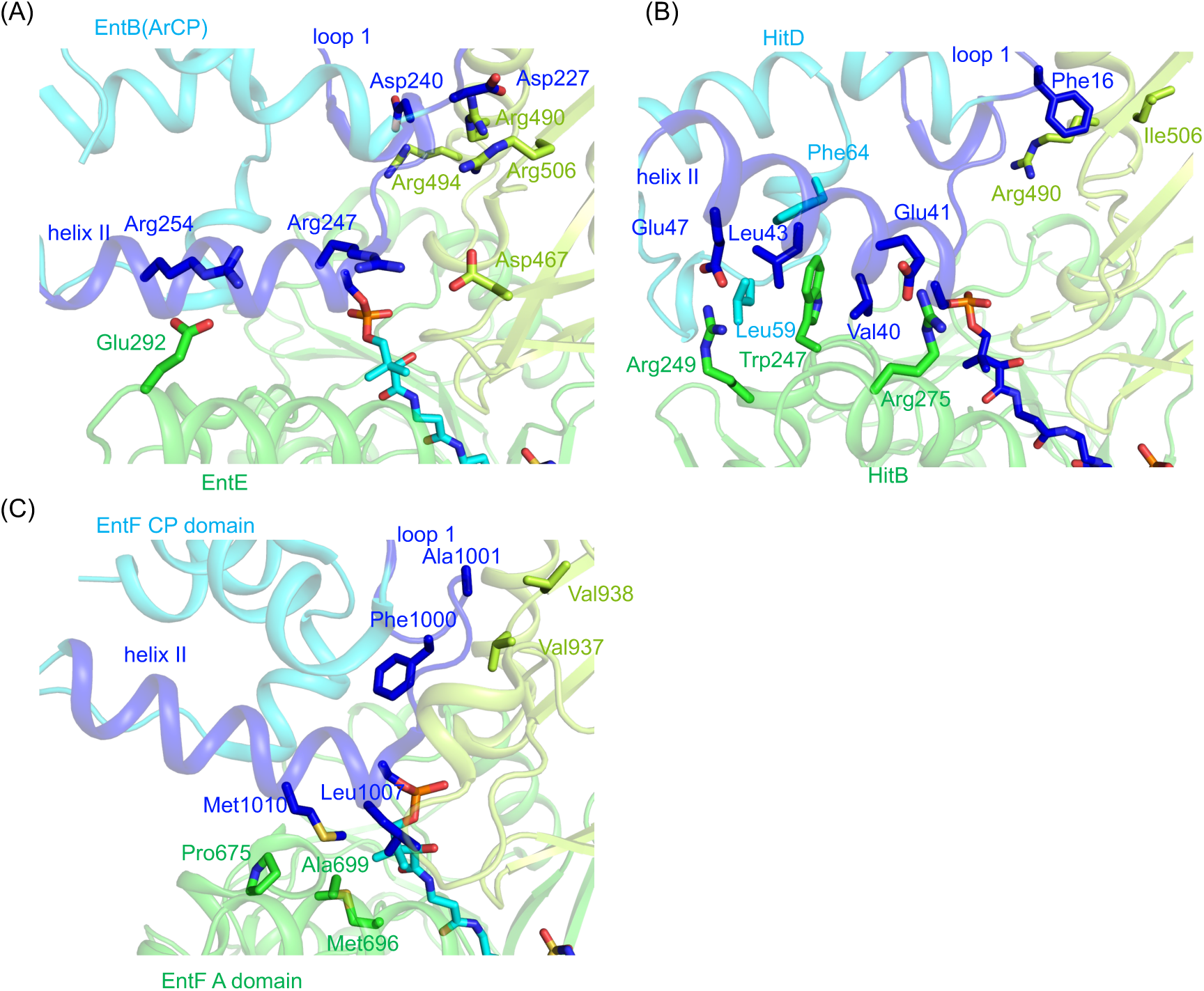
Structural comparison of the binding interface between A domain and CP. The A_core_ and A_sub_ regions of A domains are colored green and yellow-green, respectively. Loop 1 and helix II regions of CPs are colored blue, while other regions of CPs are colored cyan. (A) Binding interface in the EntE–EntB(ArCP) complex (PDB ID: 3RG2). (B) Binding interface in the HitB–HitD complex (PDB ID: 6M01). (C) Binding interface in the EntF A domain–CP complex (PDB ID: 5T3D).

**Extended Data Fig. 2.**
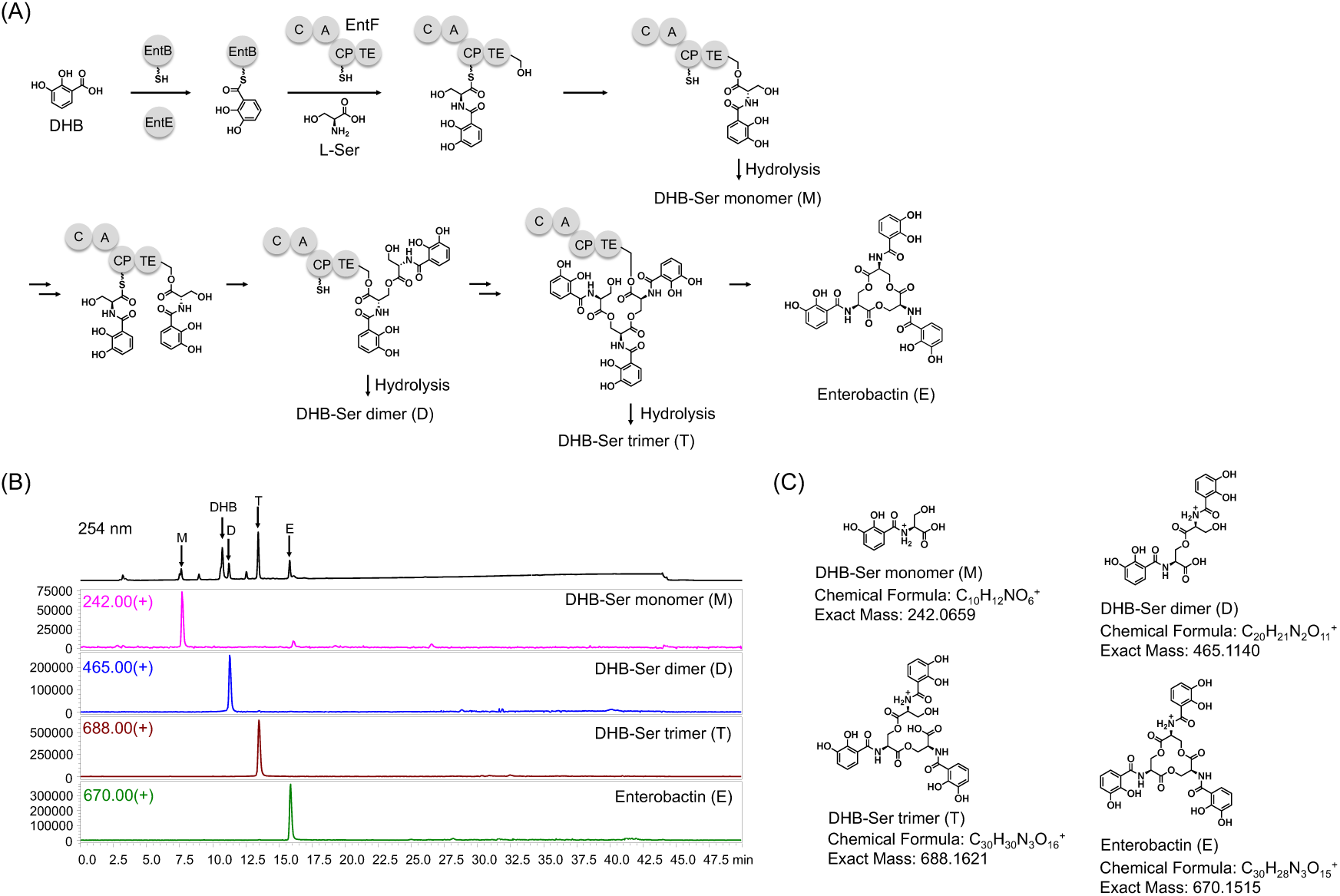
Biosynthetic pathway of the siderophore enterobactin. (A) Detailed biosynthetic pathway of enterobactin and its early-stage intermediates. EntF comprises carrier protein (CP), adenylation (A), condensation (C), and thioesterase (TE) domains. (B) HPLC and LC-MS data for the *in vitro* reaction with wtEntE, EntB, and EntF in the presence of DHB, L- serine, and ATP. (C) Structures of DHB-Ser-monomer (M), DHB-Ser-dimer (D), DHB-Ser-trimer (T), and enterobactin (E).

**Extended Data Fig. 3.**
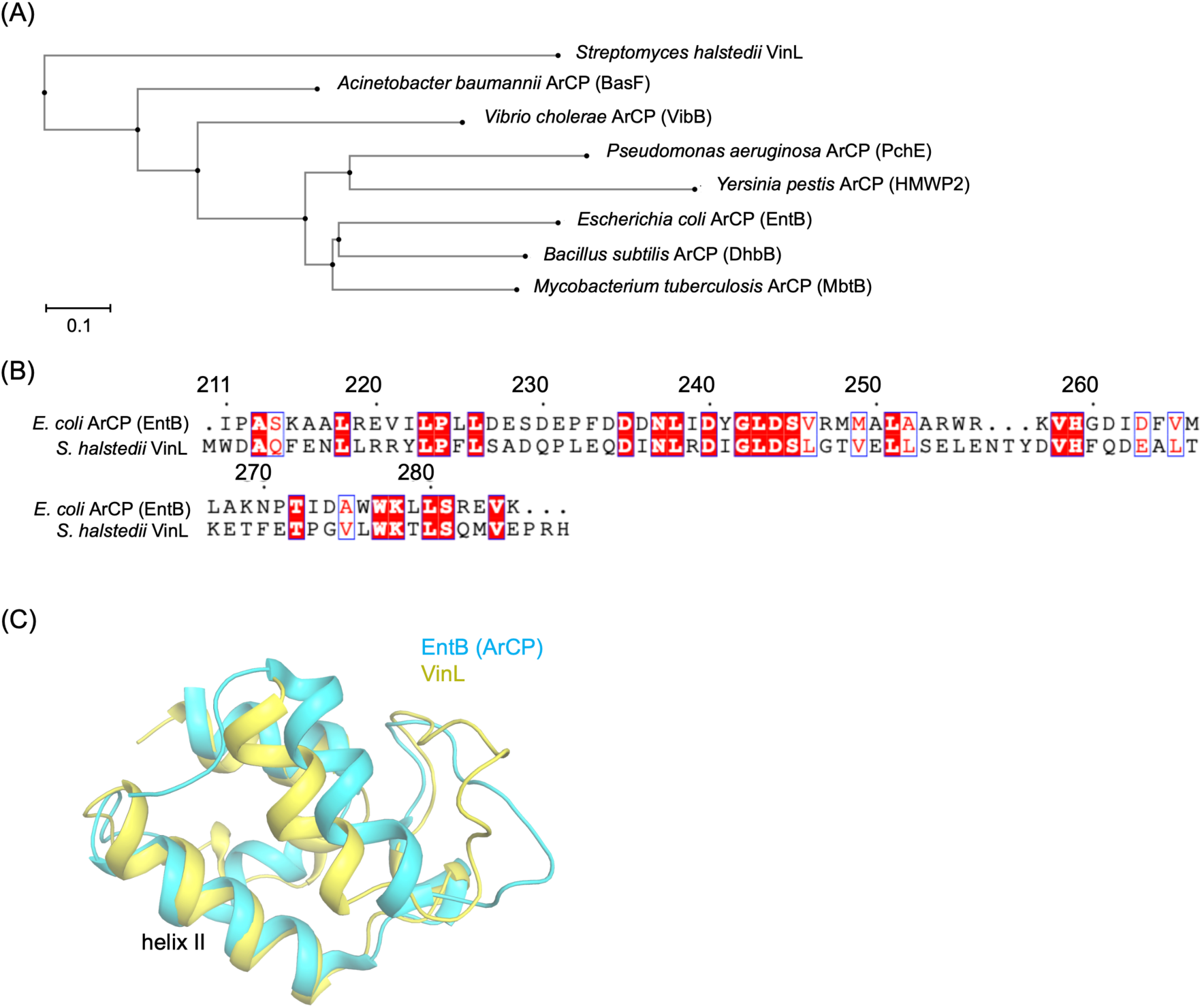
Phylogenetic and structural comparison of VinL with representative aryl carrier proteins (ArCPs). (A) Phylogenetic tree. (B) Sequence alignment between EntB(ArCP) and VinL. (C) Superimposition of EntB(ArCP) and VinL structures. EntB(ArCP) and VinL are colored cyan and yellow, respectively.

**Extended Data Fig. 4.**
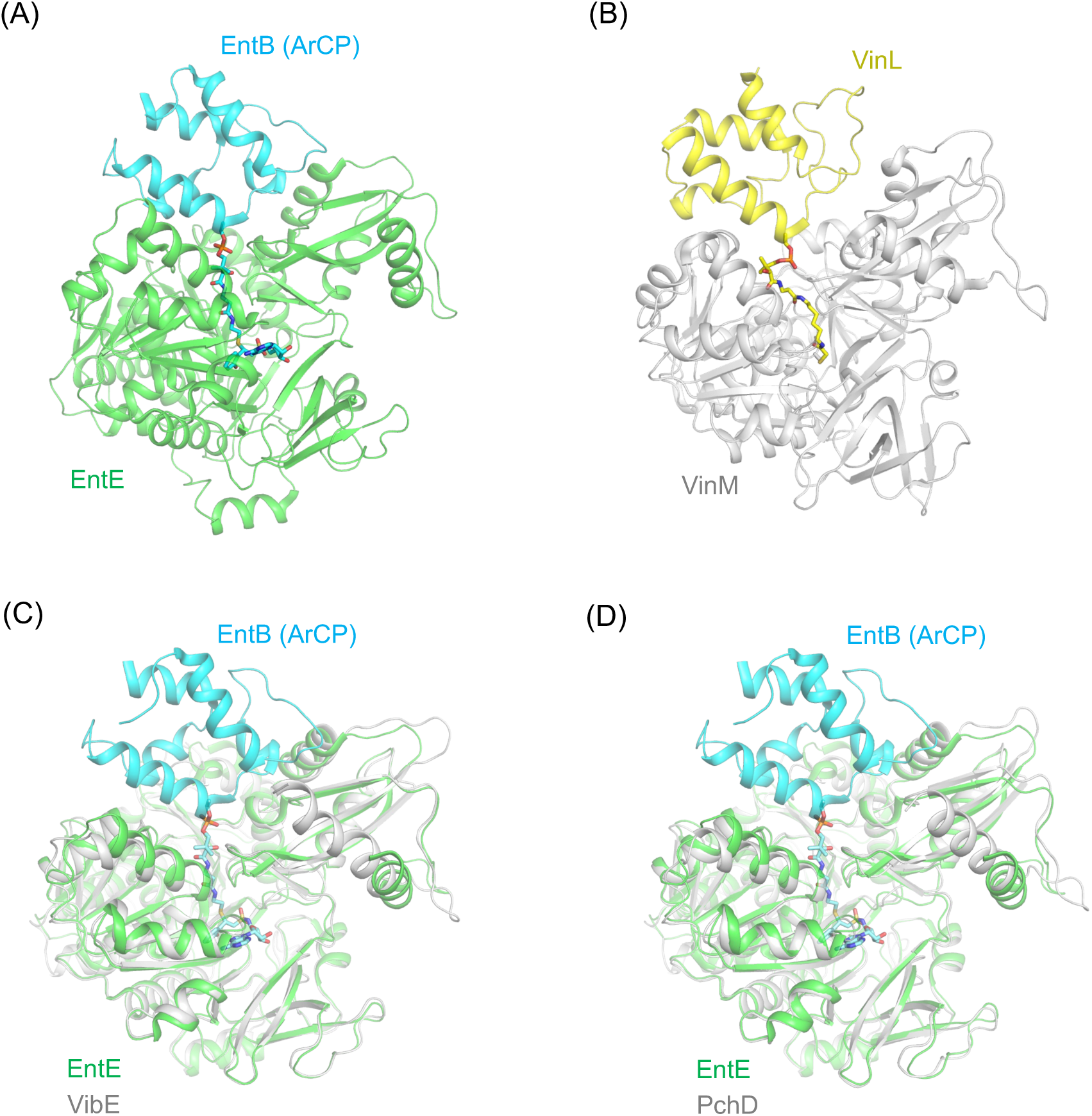
Structural comparison of cognate and modeled non-cognate adenylation–carrier protein interfaces. (A) Overall structure of the EntE–EntB(ArCP) complex (PDB ID: 3RG2). (B) Overall structure of the VinM–VinL complex (PDB ID: 8K4R). (C) Superimposition of the EntE–EntB(ArCP) complex structure and the VibE AlphaFold3-predicted structure. (D) Superimposition of the EntE–EntB(ArCP) complex structure and the PchD AlphaFold3-predicted structure. EntE, EntB(ArCP), and VinL are colored green, cyan and yellow, respectively. VinM, VibE and PchD are colored light gray.

**Extended Data Fig. 5.**
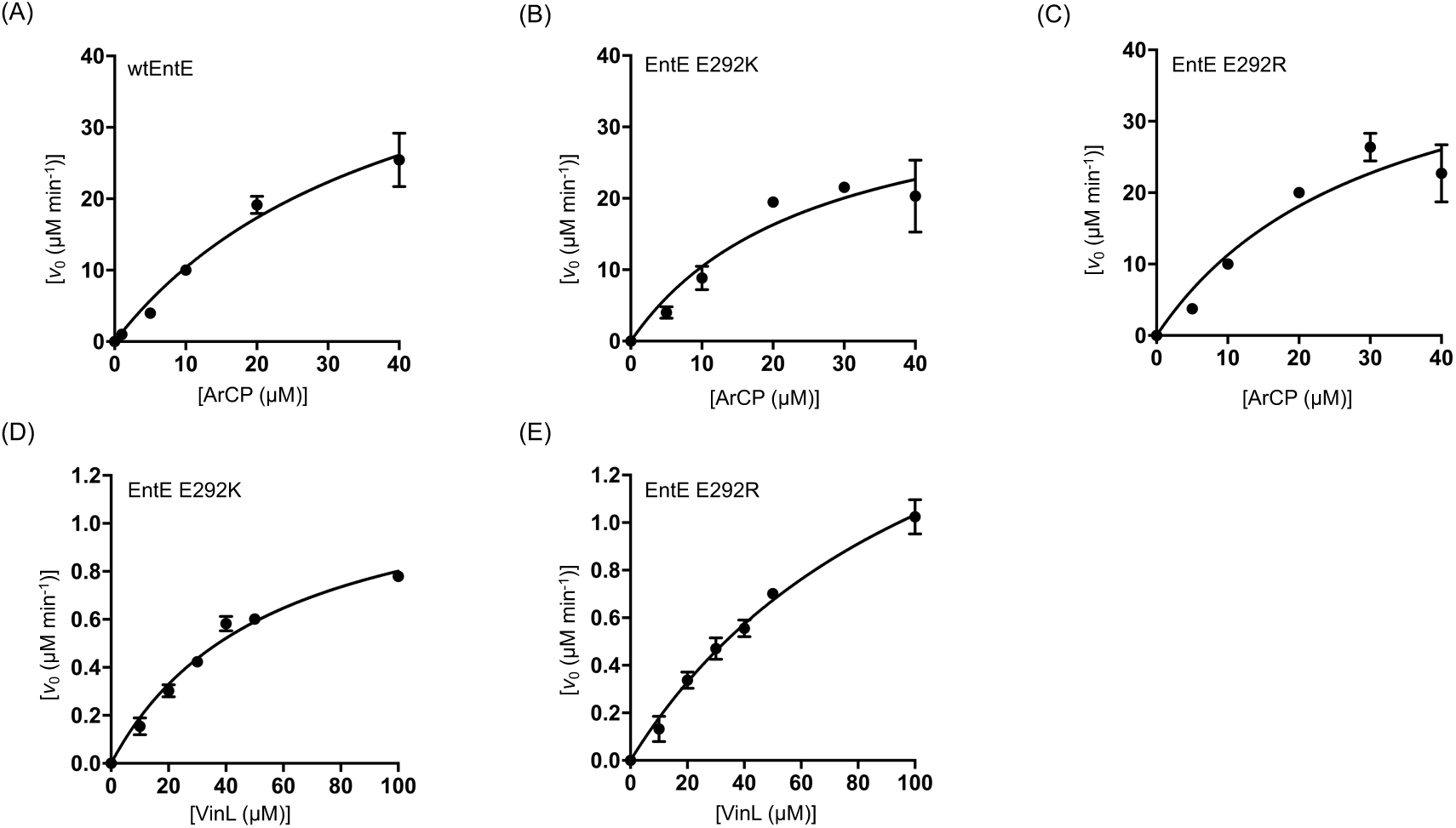
Determination of apparent kinetic constants for DHB transfer by wtEntE and EntE mutants to EntB(ArCP). (A) Each reaction contained *holo*-EntB(ArCP) (1– 40 µM), 100 nM wtEntE, 3.1 mM ATP, and 1.25 mM DHB. (B) Each reaction contained *holo*- EntB(ArCP) (5–40 µM), 100 nM EntE E292K, 3.1 mM ATP, and 1.25 mM DHB. (C) Each reaction contained *holo*-EntB(ArCP) (5–40 µM), 100 nM EntE E292R, 3.1 mM ATP, and 1.25 mM DHB. (D) Each reaction contained *holo*-VinL (10–100 µM), 200 nM EntE E292K, 3.1 mM ATP, and 1.25 mM DHB. (E) Each reaction contained *holo*-VinL (10–100 µM), 200 nM EntE E292R, 3.1 mM ATP, and 1.25 mM DHB. All reactions were performed in a buffer consisting of 50 mM HEPES–NaOH (pH 8.0), 10 mM MgCl_2_, and 5 mM DTT. Errors are shown as mean ± S.D. (*n* = 3 independent measurements).

**Extended Data Fig. 6.**
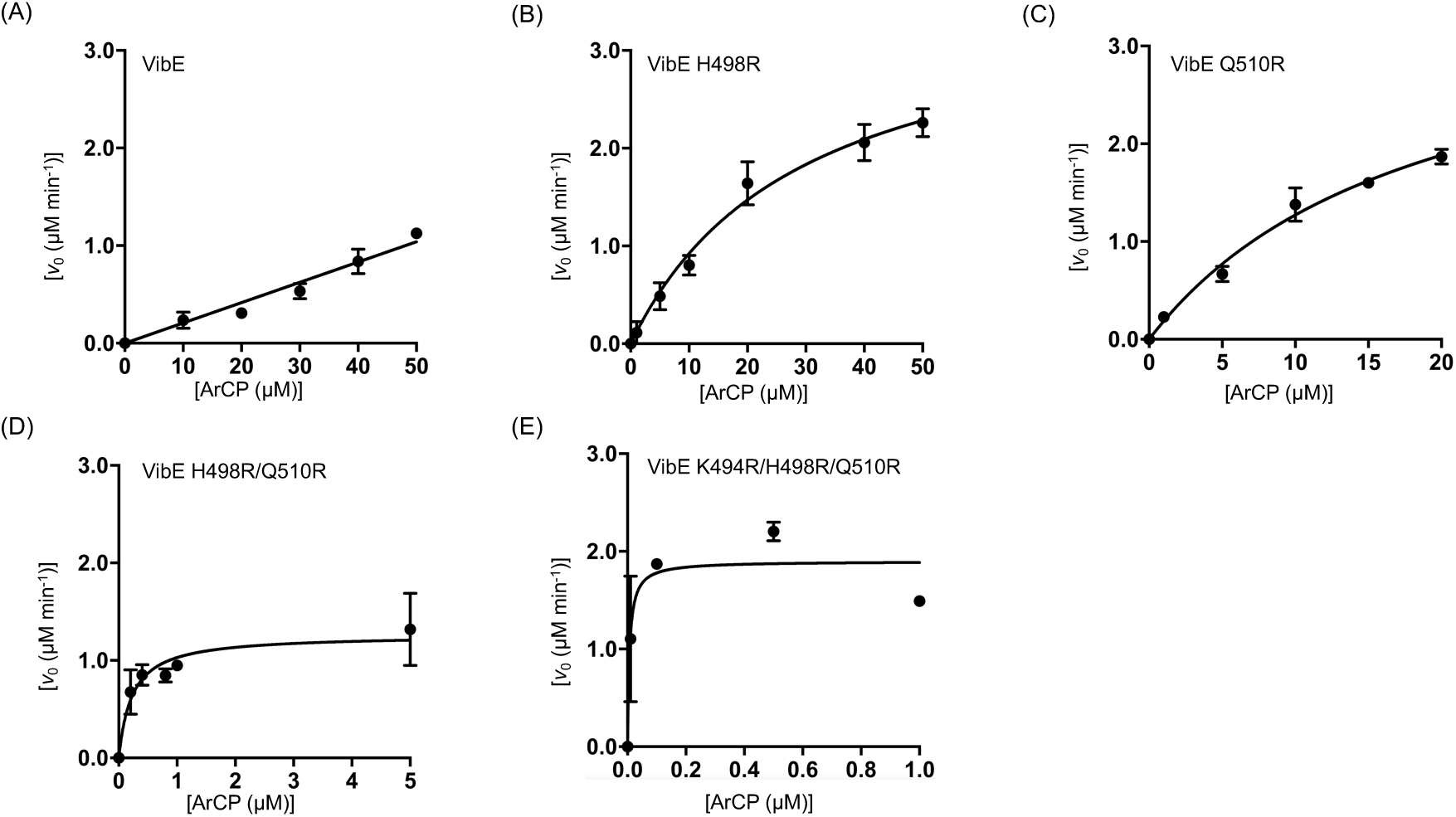
Determination of apparent kinetic constants for DHB transfer by wtVibE and VibE mutants to EntB(ArCP). (A) Each reaction contained *holo*-EntB(ArCP) (10– 50 µM), 100 nM wtVibE, 200 μM ATP, and 1 mM DHB. (B) Each reaction contained *holo*- EntB(ArCP) (1–50 µM), 100 nM VibE H498R, 200 μM ATP, and 1 mM DHB. (C) Each reaction contained *holo*-EntB(ArCP) (1–20 µM), 100 nM VibE Q510R, 200 μM ATP, and 1 mM DHB. (D) Each reaction contained *holo*-EntB(ArCP) (0.2–5 µM), 500 nM VibE H498R/Q510R, 200 μM ATP, and 1 mM DHB. (E) Each reaction contained *holo*-EntB(ArCP) (0.01–1 µM), 2 μM VibE K494R/H498R/Q510R, 200 μM ATP, and 1 mM DHB. All reactions were performed in a buffer consisting of 75 mM Tris-HCl (pH 7.5), 10 mM MgCl_2_, and 5 mM DTT. Errors are shown as mean ± S.D. (*n* = 2–3 independent measurements).

**Extended Data Fig. 7.**
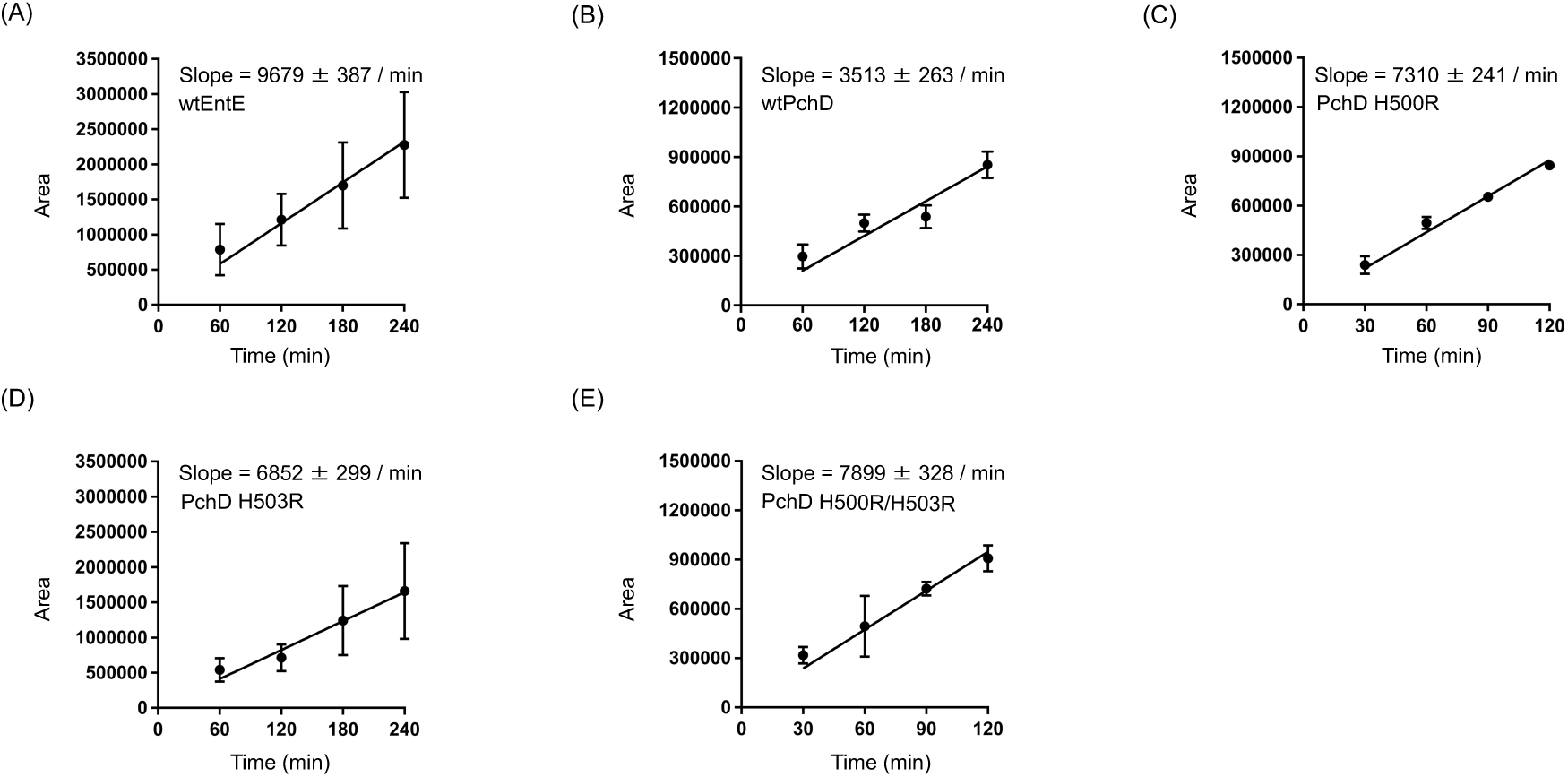
Initial velocities for Sal-NSPD production by wtEntE, wtPchD, and PchD mutants. (A) Each reaction contained *holo*-EntB (10 µM), 1 µM wtEntE, 1 µM VibH, 5 mM ATP, 1 mM Sal, and 5 mM NSPD. (B) Each reaction contained *holo*-EntB (10 µM), 1 µM wtPchD, 1 µM VibH, 5 mM ATP, 1 mM Sal, and 5 mM NSPD. (C) Each reaction contained *holo*- EntB (10 µM), 1 µM PchD Q500R, 1 µM VibH, 5 mM ATP, 1 mM Sal, and 5 mM NSPD. (D) Each reaction contained *holo*-EntB (10 µM), 1 µM PchD H503R, 1 µM VibH, 5 mM ATP, 1 mM Sal, and 5 mM NSPD. (E) Each reaction contained *holo*-EntB (10 µM), 1 µM PchD Q500R/H503R, 1 µM VibH, 5 mM ATP, 1 mM Sal, and 5 mM NSPD. All reactions were performed in a buffer consisting of 50 mM HEPES–NaOH (pH 8.0), 10 mM MgCl_2_, and 5 mM DTT. Errors are shown as mean ± S.D. (*n* = 3 independent measurements).

**Extended Data Fig. 8.**
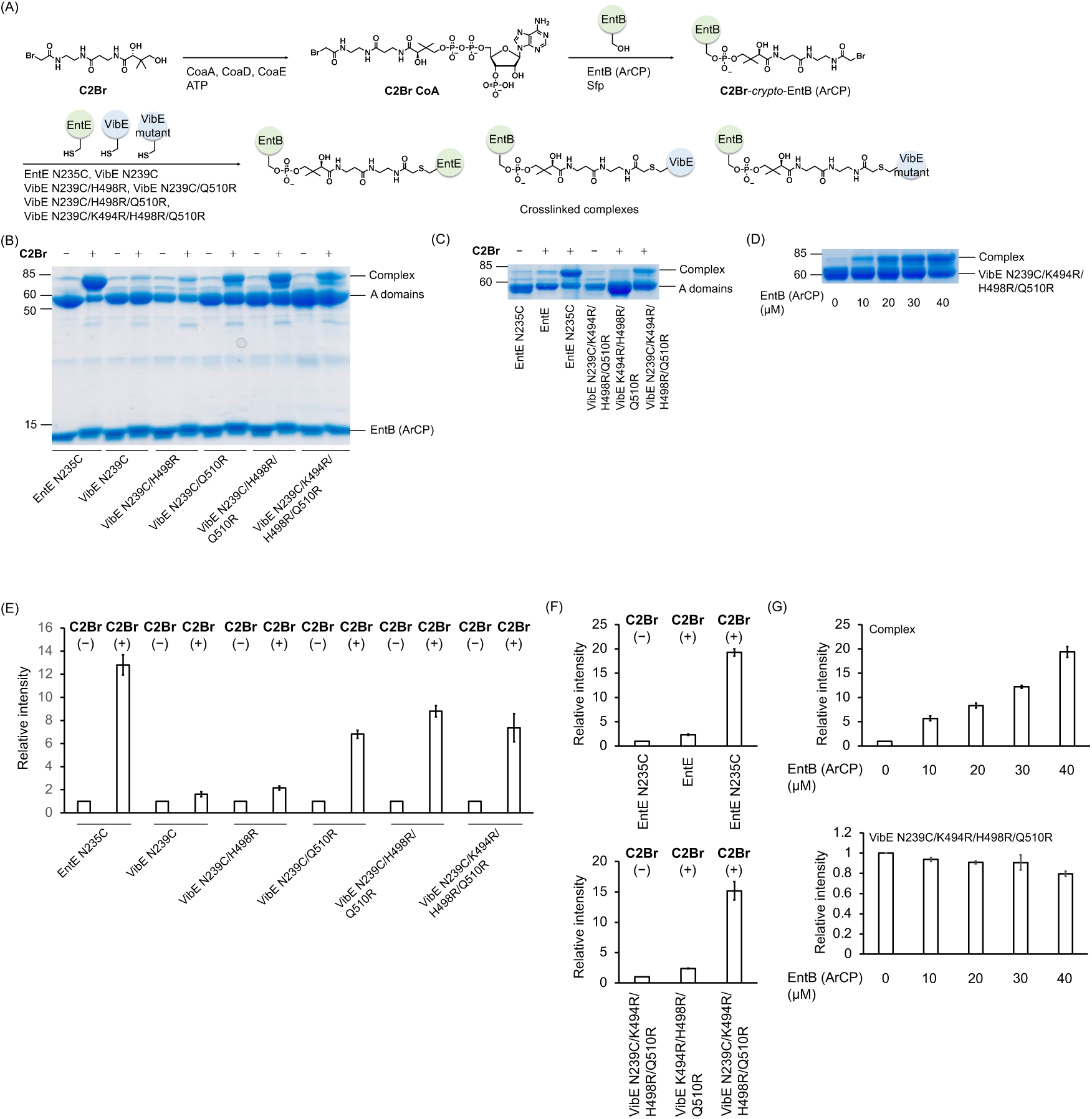
Crosslinking strategy. (A) Schematic of the three-step, one-pot crosslinking reaction. Crosslinker **C2Br** was converted to the corresponding coenzyme A analogue, **C2Br CoA**, which was then loaded in situ onto the conserved serine residue of EntB(ArCP) (Ser245) by Sfp, a 4’-phosphopantetheinyltransferase (PPTase). After loading of EntB(ArCP) was complete, the samples were reacted with EntE N235C, VibE N239C, VibE N239C/H498R, VibE N239C/Q510R, VibE N239C/H498R/Q510R, and VibE N239C/K494R/H498R/Q510R to generate crosslinked complexes. (B) Crosslinking reactions of EntB(ArCP) and adenylation enzymes. (C) Crosslinking efficiency and validation of cysteine reactivity. (D) Concentration dependence of crosslinking on EntB(ArCP) concentration. (E) Quantification of crosslinked complex bands in (B). Intensities of the complex bands were quantified and expressed relative to the corresponding DMSO controls. (F) Quantification of crosslinked complex bands in (C). Intensities of the complex bands were quantified and expressed relative to the corresponding DMSO controls. (G) Quantification of the crosslinked complex and VibE N239C/K494R/H498R/Q510R bands in (D). Intensities of the crosslinked complex and VibE N239C/K494R/H498R/Q510R bands were quantified and expressed relative to the condition without EntB(ArCP). Data are shown as mean ± S.D. (n = 2 independent measurements). All one-pot crosslinking reactions included treatment with EntB(ArCP), **C2Br**, CoaA, CoaD, CoaE, and Sfp for 1 h at 37 ℃. **C2Br**-*crypto*-EntB(ArCP) was then incubated with 5 μM adenylation enzymes for 16 h at 37 ℃. Uncropped gels for the images in (B–D) are provided in Supplementary Fig. S21.

**Extended Data Fig. 9.**
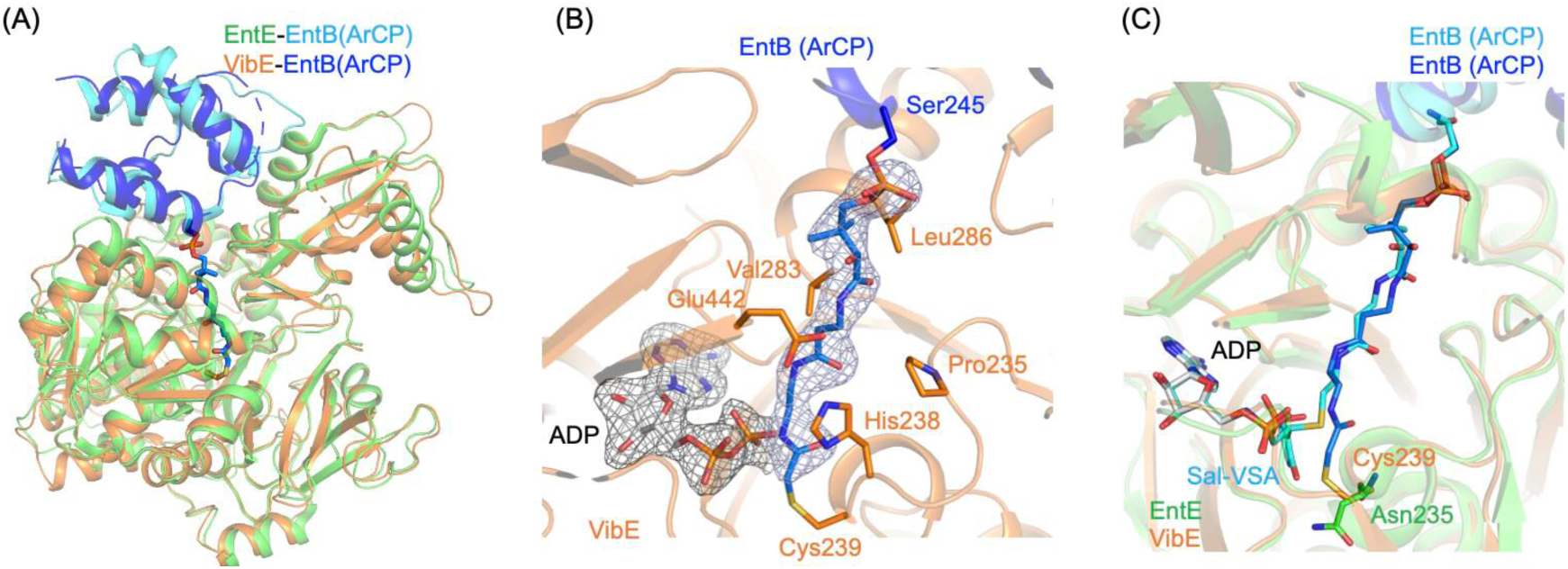
Structural comparison of the VibE–EntB(ArCP) complex with the EntE–EntB(ArCP) complex. (A) Superimposition of the VibE–EntB(ArCP) complex and the EntE–EntB(ArCP) complex (PDB ID: 3RG2). VibE adopts a thioester-forming conformation, similar to that observed for EntE in the EntE–EntB(ArCP) complex structure. VibE and EntB(ArCP) in the VibE–EntB(ArCP) complex are colored orange and blue, respectively. EntE and EntB(ArCP) in the EntE–EntB(ArCP) complex are colored green and cyan, respectively. (B) The pantetheine-binding tunnel in the VibE–EntB(ArCP) complex structure. Electron density corresponding to the phosphopantetheine linker derived from **C2Br** was observed in the pantetheine-binding tunnel. The phosphopantetheine linker covalently connects Ser245 of EntB(ArCP) and the mutated Cys239 of VibE. Additional electron density was observed at the ATP-binding site of VibE. This electron density might be a mixture of ADP and AMP molecules, potentially derived from ATP used during enzymatic synthesis of *crypto*-EntB(ArCP), as previously proposed for the HitB–HitD complex structure.^10^ An ADP molecule is shown as light gray sticks. The phosphopantetheine linker is shown as blue sticks. Polder maps for the phosphopantetheine linker moiety (contoured at 5σ) and the ADP molecule (contoured at 5σ) are shown as blue and gray meshes. (C) Structural comparison of the pantetheine-binding tunnels of the VibE–EntB(ArCP) complex and the EntE–EntB(ArCP) complex.

**Extended Data Fig. 10.**
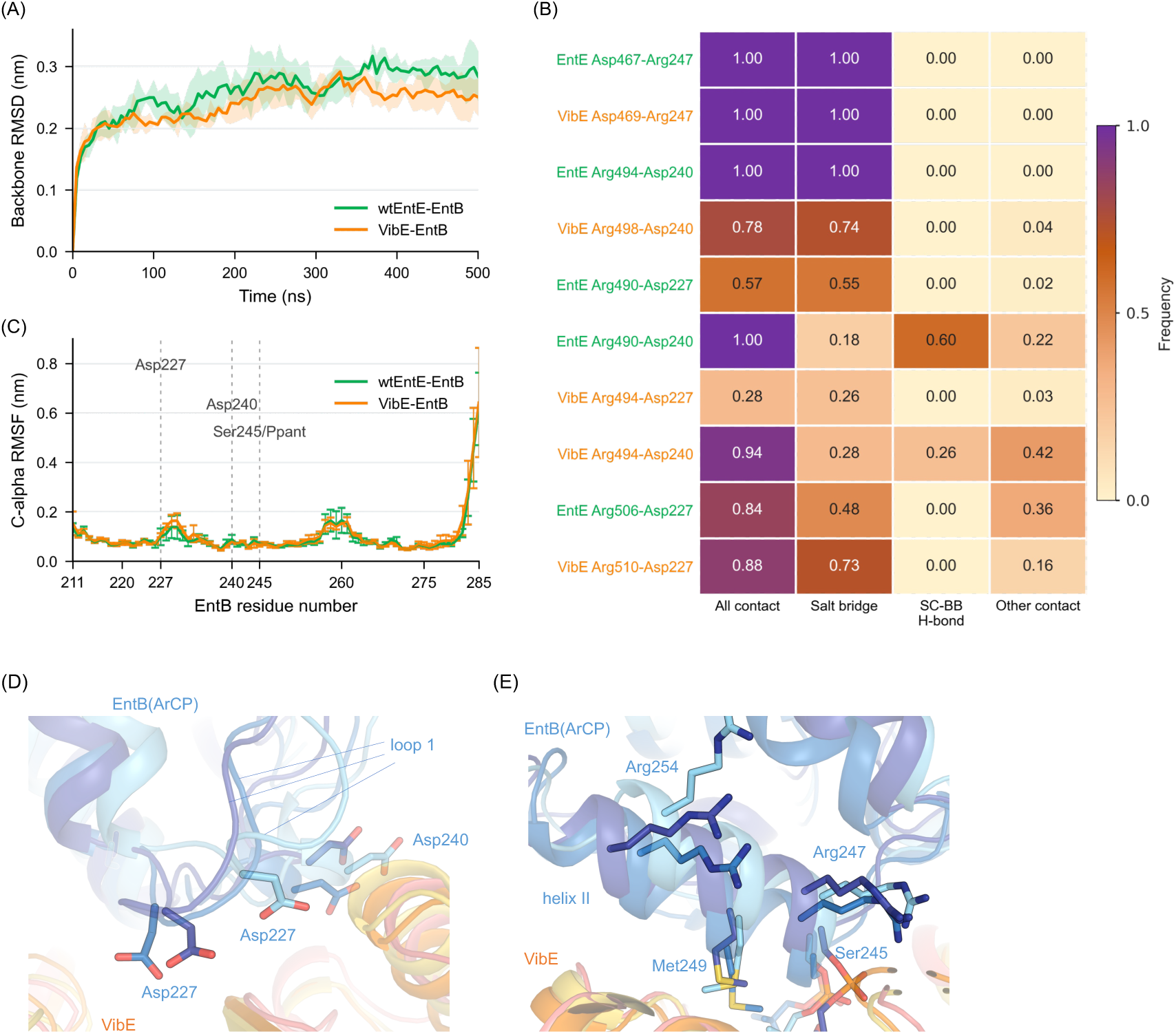
Structural dynamics of A–CP complexes during MD simulations. Structural stability and interaction frequencies of the complexes during MD simulations. (A) Backbone RMSD profiles of the wtEntE–EntB(ArCP) and VibE K494R/H498R/Q510R– EntB(ArCP) complexes from triplicate 500-ns production simulations. Lines and shaded regions indicate the mean and S.D. across triplicate simulations, respectively. RMSD values were calculated after backbone fitting of each complex and are shown with 5-ns binning. (B) Frequency heatmap of selected inter-protein interactions around EntB Asp227 and Asp240 across triplicate 500-ns simulations. Interaction frequencies indicate the proportion of analyzed trajectory frames, sampled every 10 ps, in which each interaction was present. “All contact” was defined using a residue-pair heavy-atom minimum-distance cutoff of ≤ 0.45 nm, “Salt bridge” using a charged- atom minimum-distance cutoff of ≤ 0.40 nm, and “SC–BB H-bond” as an Arg side-chain to Asp/Glu backbone hydrogen bond. “Other contact” indicates contact frames without either a salt bridge or an SC–BB H-bond. (C) Cα RMSF of EntB(ArCP) calculated from trajectories sampled every 100 ps after fitting to EntB Cα atoms. Lines and error bars indicate the mean and S.D. across triplicate simulations, respectively. EntB loop 1, including Asp227 and Asp240, and the phosphopantetheinylated Ser245 are indicated. (D,E) Superimposition of three representative conformations of the VibE–EntB(ArCP) complex during MD simulations. (D) Close-up view around EntB loop 1 at the VibE–EntB(ArCP) interface, showing conformational diversity of the loop containing Asp227 and Asp240. EntB Asp227 and Asp240 are shown as sticks. (E) Close- up view around the VibE A_core_-facing interface near EntB Arg254, showing that Arg254 is oriented away from the VibE A_core_ region and does not form an EntE-like salt bridge. EntB Met249, Arg247, and phosphopantetheinylated Ser245 are also shown as sticks.

