## Supplementary Notes, and Supplementary Figure 1-21, Supplementary Table 1, and Supplementary References for "Reprogramming protein interfaces between adenylation and carrier protein domains in nonribosomal peptide synthetases"

### Table of contents


**Supplementary Notes**……………………………………………………………………………………………3
**Supplementary Fig. 1.** SDS-PAGE analysis of purified EntE E292K…………………………………………………..5
**Supplementary Fig. 2.** SDS-PAGE analysis of purified EntE E292R……………….……………………………….6
**Supplementary Fig. 3.** SDS-PAGE analysis of purified VibE H498R…………………………...………………..7
**Supplementary Fig. 4.** SDS-PAGE analysis of purified VibE Q510R…………………………………………… …..8
**Supplementary Fig. 5.** SDS-PAGE analysis of purified VibE H498R/Q510R……………………………………….....9
**Supplementary Fig. 6.** SDS-PAGE analysis of purified VibE K494R/H498R/Q510R……………………………...10
**Supplementary Fig. 7.** SDS-PAGE analysis of purified PchD Q500R……………………………………...………11
**Supplementary Fig. 8.** SDS-PAGE analysis of purified PchD H503R…………………………………………… …12
**Supplementary Fig. 9.** SDS-PAGE analysis of purified PchD Q500R/H503R………………………..……………13
**Supplementary Fig. 10.** SDS-PAGE analysis of purified VibE N239C…………………….……………………14
**Supplementary Fig. 11.** SDS-PAGE analysis of purified VibE N239C/H498R……………………………...…..15
**Supplementary Fig. 12.** SDS-PAGE analysis of purified VibE N239C/Q510R……………………...……………..16
**Supplementary Fig. 13.** SDS-PAGE analysis of purified VibE N239C/H498R/Q510R………...…………………17
**Supplementary Fig. 14.** SDS-PAGE analysis of purified VibE N239C/K494R/H498R/Q510R……….…………..18
**Supplementary Fig. 15.** Expanded MALDI-TOF mass spectra of DHB loading onto EntB(ArCP) by wtEntE……….19
**Supplementary Fig. 16.** Expanded MALDI-TOF mass spectra of DHB loading onto EntB(ArCP) by EntE E292K….20
**Supplementary Fig. 17.** Expanded MALDI-TOF mass spectra of DHB loading onto EntB(ArCP) by EntE E292R….21
**Supplementary Fig. 18.** Expanded MALDI-TOF mass spectra of DHB loading onto VinL by wtEntE…………...…..22
**Supplementary Fig. 19.** Expanded MALDI-TOF mass spectra of DHB loading onto VinL by EntE E292K…………23
**Supplementary Fig. 20.** Expanded MALDI-TOF mass spectra of DHB loading onto VinL by EntE E292R……...…..24
**Supplementary Fig. 21.** Uncropped SDS-PAGE gels for crosslinking reactions…………………...……………….25
**Supplementary Table 1.** X-ray data collection and refinement statistics……………………………………...…27
**Supplementary References**……………………………………………………………………...…………28

**Supplementary Notes**

Over the past decade, pantetheinamide crosslinkers have been developed and applied to the structural analysis of carrier proteins (CPs) in complex with partner domains from fatty acid, polyketide, and nonribosomal peptide biosynthesis.[1,2] For example, sulfonyl 3-alkynyl pantetheinamides have been used to capture transient dehydratase–CP complexes through covalent linkage to the catalytic His residue of the dehydratase.[3,4] Likewise, a chloroacryl pantetheinamide was used to exploit the nucleophilicity of the active-site Cys residue of ketosynthase,[5] and a C5-phenylsulfonyl pantetheinamide efficiently generated an intramolecular epimerase–CP crosslinked complex by reacting with the catalytic His residue of the epimerase.[6] More recently, we determined the structure of an acyltransferase–CP complex using the bromoacetyl pantetheinamide **C2Br** in combination with a Cys mutation of acyltransferase (Extended Data Fig. 8).[7] We further extended this **C2Br**-based strategy to structurally characterize adenylation (A)–CP crosslinked complexes, including the HitB–HitD complex generated by introducing a Cys mutation at Asp221 of HitB and the VinM–VinL complex generated by introducing a Cys mutation at Asp223 of VinM.[8,9] Collectively, these studies highlight the importance of expanding site-selective crosslinking strategies to enable capture of non-native protein–protein interaction pairs. The Asn235 residue of EntE and the corresponding Asn239 residue of VibE, which align with Asp221 of HitB and Asp223 of VinM, are positioned in the substrate-binding pocket and engage in hydrogen-bonding interactions with the 2- and 3-hydroxyl groups of 2,3-dihydroxybenzoic acid (DHB).[8,9,10,] Thus, we incorporated a Cys mutation at Asn239 of VibE, anticipating that this residue would serve as a suitable nucleophile for reaction with the bromoacetyl group of **C2Br**. Recombinant VibE N239C, VibE N239C/H498R, VibE N239C/Q510R, VibE N239C/H498R/Q510R, and VibE N239C/K494R/H498R/Q510R mutants were overproduced and purified as *C*-terminally $His_6$-tagged constructs (Supplementary Fig. S10–S14). For comparative activity studies, recombinant EntE N235C was prepared according to the procedure reported previously.[10]

We loaded **C2Br** onto the Ser245 residue of the *apo*-ArCP of EntB in an one-pot chemo-enzymatic strategy using CoaA, CoaD, CoaE, and Sfp (Extended Data Fig. 8).[10] Recombinant CoaA, CoaD, CoaE, and Sfp were overproduced and purified, as described previously.[11] EntE N235C, VibE N239C, VibE N239C/H498R, VibE N239C/Q510R, VibE N239C/H498R/Q510R, and VibE N239C/K494R/H498R/Q510R mutants were then reacted with the modified EntB(ArCP) (Extended Data Fig. 8). As shown in Extended Data Fig. 8, crosslinking was observed for EntE N235C, VibE N239C/H498R, VibE N239C/Q510R, VibE N239C/H498R/Q510R, and VibE N239C/K494R/H498R/Q510R, as indicated by a gel shift from ~60 kDa to ~85 kDa on SDS-PAGE. In contrast, no crosslinked species was detected for VibE N239C in the presence of *crypto*-EntB(ArCP) (Extended Data Fig. 8). This result is fully consistent with our earlier biochemical data indicating that wtVibE does not productively interact with EntB(ArCP), and suggests that the wild-type interface is insufficient to support stable complex capture under these conditions (Fig. 3H and 4A bottom). On the basis of these results, we selected VibE N239C/K494R/H498R/Q510R for subsequent structural and functional analyses. Among the variants examined, this mutant exhibited the lowest apparent $K_m$ for EntB(ArCP) and supported the highest level of enterobactin production. Crosslinking of VibE N239C/K494R/H498R/Q510R with **C2Br**-loaded EntB(ArCP) was observed as a clear band shift from ~60 kDa to ~85 kDa on SDS-PAGE, indicating formation of the covalently captured CP–A domain complex (Extended Data Fig. 8). By contrast, no crosslinked species was detected for the corresponding Cys-free mutant VibE K494R/H498R/Q510R under the same conditions, indicating that complex formation depends on the introduced Cys residue (Extended Data Fig. 8). Although the crosslinking efficiency was lower than that observed for the control EntE N235C–EntB(ArCP) pair, it was sufficient to support subsequent structural studies, as the abundance of the crosslinked VibE complex increased in an EntB(ArCP)-dependent manner (Extended Data Fig. 8). Thus, the site-selective crosslinking experiments provide structural support for the conclusion that interface reprogramming promotes formation of a productive non-cognate A–CP complex. The engineered VibE–EntB complex could be covalently captured in a Cys-dependent manner, whereas VibE N239C failed to form a crosslinked species. Together with the concentration dependence of complex formation, these data indicate that interface reprogramming promotes formation of a structurally accessible and functionally productive non-cognate A–CP complex.

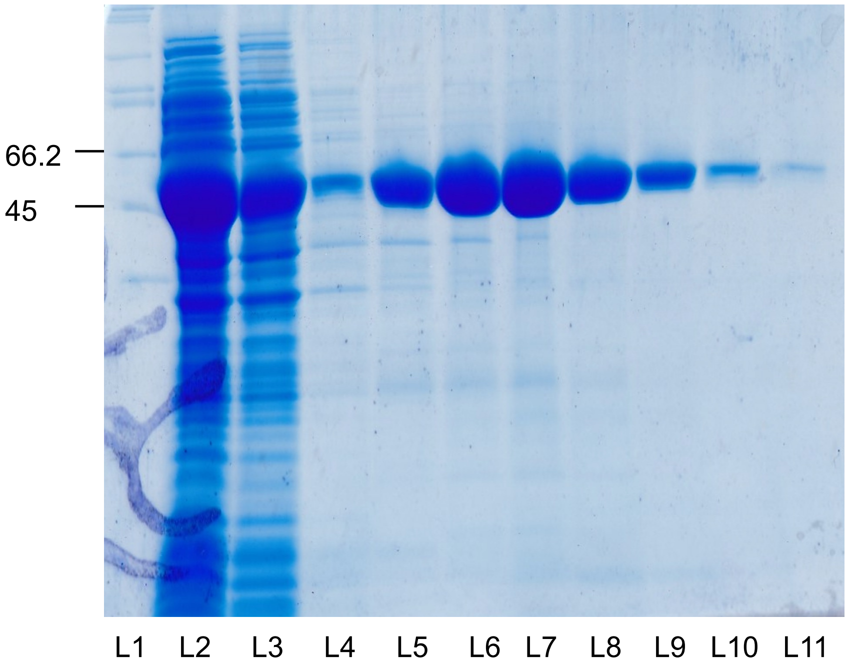


**Supplementary Fig. 1. Full gel depicting the purification of the mutant EntE E292K.** Gel lanes depict fractions taken during Ni-sepharose chromatography (Nacalai Tesque, Inc.) and are as follows: L1 = protein marker, L2 = flow through, L3= wash, L4 = 10 mM imidazole wash, L5 = 20 mM imidazole wash, L6 = 40 mM imidazole wash, L7 = 80 mM imidazole wash, L8 = 100 mM imidazole wash, L9 = 150 mM imidazole wash, L10 = 200 mM imidazole wash, and L11 = 400 mM imidazole wash. The target protein was collected and used from L5–L11. The gel was stained with Coomassie Brilliant Blue (CBB).

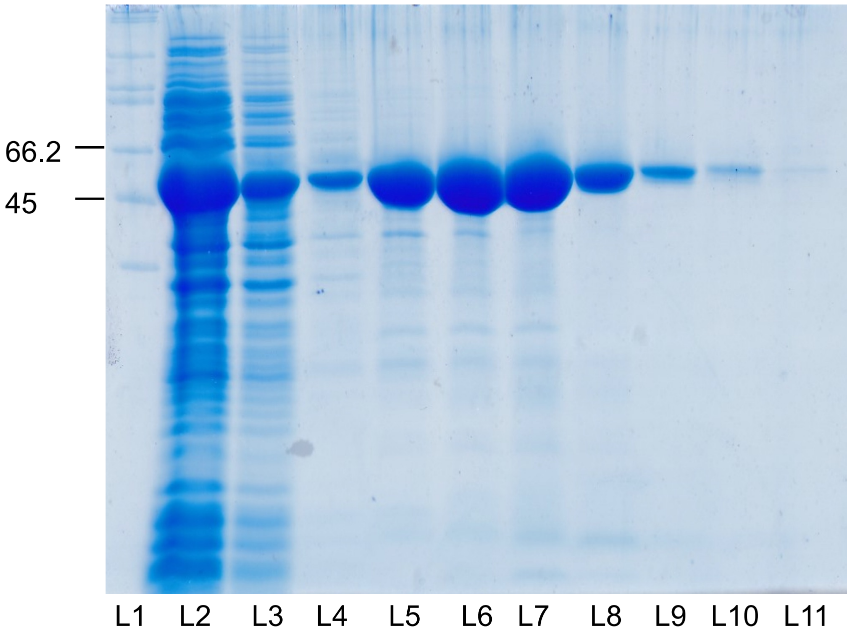


**Supplementary Fig. 2. Full gel depicting the purification of the mutant EntE E292R.** Gel lanes depict fractions taken during Ni-sepharose chromatography (Nacalai Tesque, Inc.) and are as follows: L1 = protein marker, L2 = flow through, L3= wash, L4 = 10 mM imidazole wash, L5 = 20 mM imidazole wash, L6 = 40 mM imidazole wash, L7 = 80 mM imidazole wash, L8 = 100 mM imidazole wash, L9 = 150 mM imidazole wash, L10 = 200 mM imidazole wash, and L11 = 400 mM imidazole wash. The target protein was collected and used from L5–L11. The gel was stained with Coomassie Brilliant Blue (CBB).

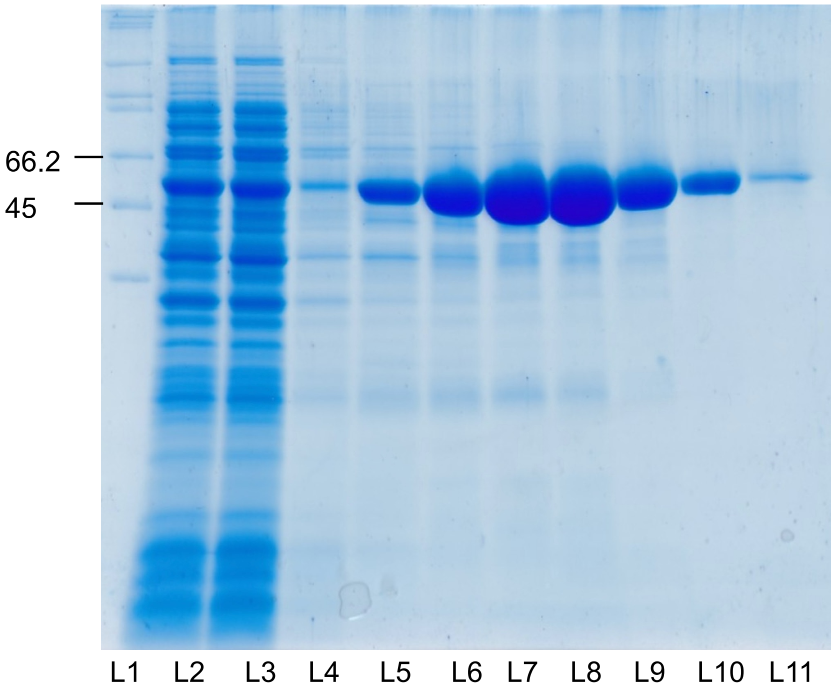


**Supplementary Fig. 3. Full gel depicting the purification of the mutant VibE H498R.** Gel lanes depict fractions taken during Ni-sepharose chromatography (Nacalai Tesque, Inc.) and are as follows: L1 = protein marker, L2 = flow through, L3= wash, L4 = 10 mM imidazole wash, L5 = 20 mM imidazole wash, L6 = 40 mM imidazole wash, L7 = 80 mM imidazole wash, L8 = 100 mM imidazole wash, L9 = 150 mM imidazole wash, L10 = 200 mM imidazole wash, and L11 = 400 mM imidazole wash. The target protein was collected and used from L5–L11. The gel was stained with Coomassie Brilliant Blue (CBB).

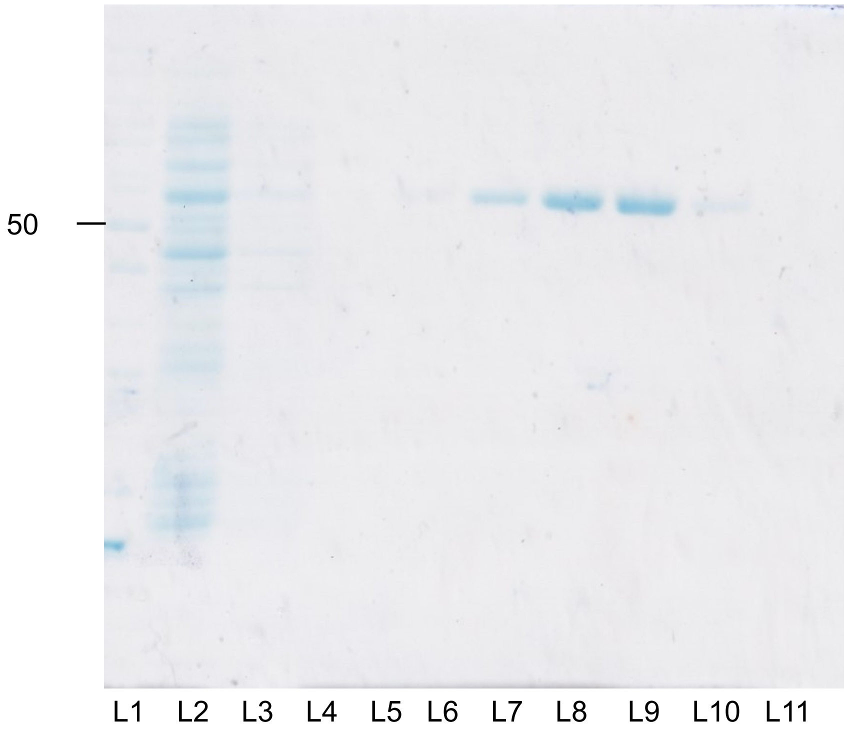


**Supplementary Fig. 4. Full gel depicting the purification of the mutant VibE Q510R.** Gel lanes depict fractions taken during Ni-sepharose chromatography (Nacalai Tesque, Inc.) and are as follows: L1 = protein marker, L2 = flow through, L3= wash, L4 = 10 mM imidazole wash, L5 = 20 mM imidazole wash, L6 = 40 mM imidazole wash, L7 = 80 mM imidazole wash, L8 = 100 mM imidazole wash, L9 = 150 mM imidazole wash, L10 = 200 mM imidazole wash, and L11 = 400 mM imidazole wash. The target protein was collected and used from L6–L10. The gel was stained with Coomassie Brilliant Blue (CBB).

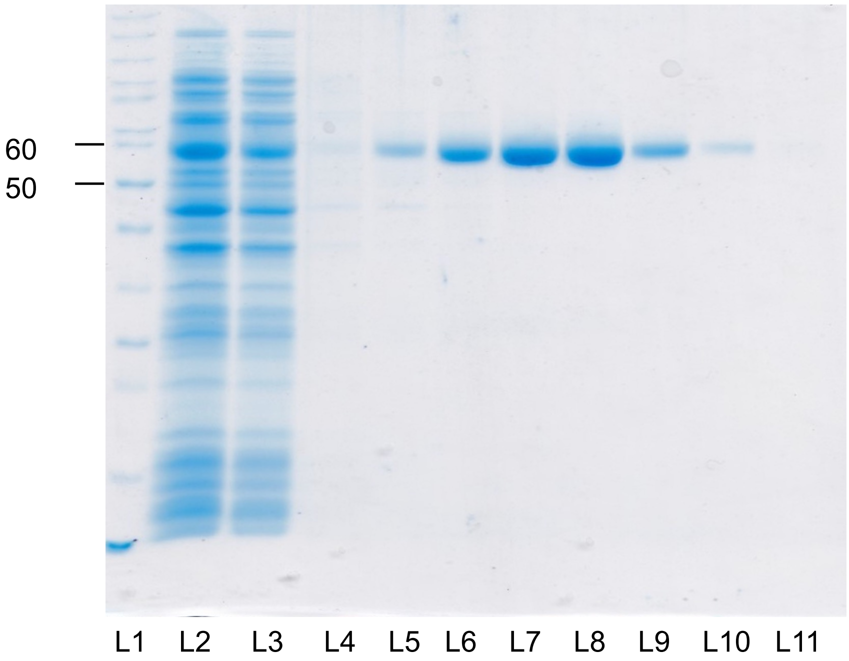


**Supplementary Fig. 5. Full gel depicting the purification of the mutant VibE H498R/Q510R.** Gel lanes depict fractions taken during Ni-sepharose chromatography (Nacalai Tesque, Inc.) and are as follows: L1 = protein marker, L2 = flow through, L3= wash, L4 = 10 mM imidazole wash, L5 = 20 mM imidazole wash, L6 = 40 mM imidazole wash, L7 = 80 mM imidazole wash, L8 = 100 mM imidazole wash, L9 = 150 mM imidazole wash, L10 = 200 mM imidazole wash, and L11 = 400 mM imidazole wash. The target protein was collected and used from L5–L10. The gel was stained with Coomassie Brilliant Blue (CBB).

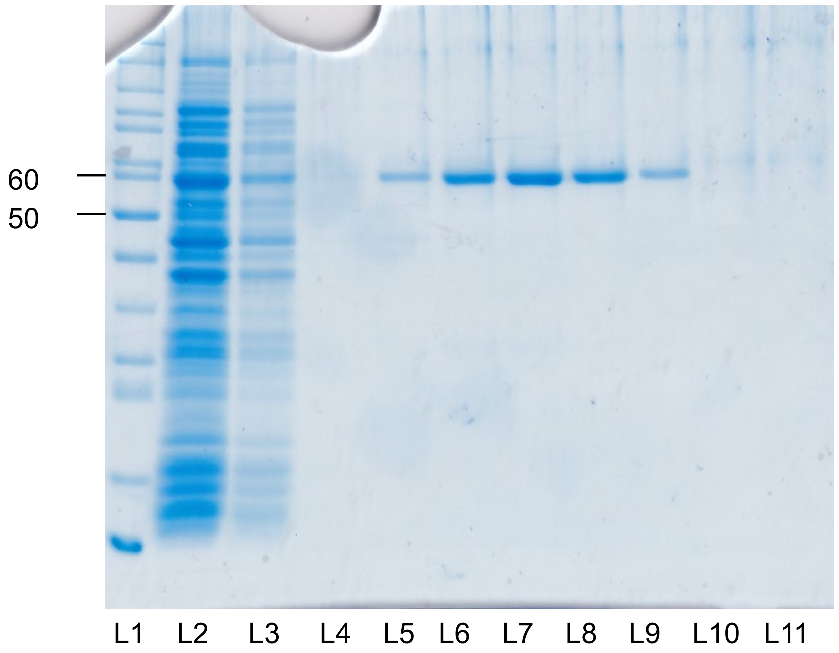


**Supplementary Fig. 6. Full gel depicting the purification of the mutant VibE K494R/H498R/Q510R.** Gel lanes depict fractions taken during Ni-sepharose chromatography (Nacalai Tesque, Inc.) and are as follows: L1 = protein marker, L2 = flow through, L3= wash, L4 = 10 mM imidazole wash, L5 = 20 mM imidazole wash, L6 = 40 mM imidazole wash, L7 = 80 mM imidazole wash, L8 = 100 mM imidazole wash, L9 = 150 mM imidazole wash, L10 = 200 mM imidazole wash, and L11 = 400 mM imidazole wash. The target protein was collected and used from L5–L9. The gel was stained with Coomassie Brilliant Blue (CBB).

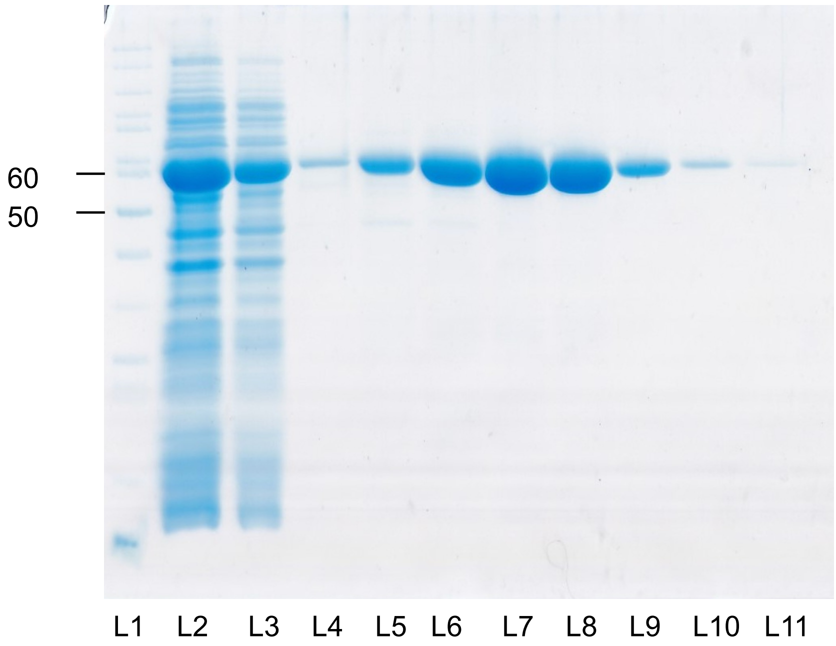


**Supplementary Fig. 7. Full gel depicting the purification of the mutant PchD Q500R.** Gel lanes depict fractions taken during Ni-sepharose chromatography (Nacalai Tesque, Inc.) and are as follows: L1 = protein marker, L2 = flow through, L3= wash, L4 = 10 mM imidazole wash, L5 = 20 mM imidazole wash, L6 = 40 mM imidazole wash, L7 = 80 mM imidazole wash, L8 = 100 mM imidazole wash, L9 = 150 mM imidazole wash, L10 = 200 mM imidazole wash, and L11 = 400 mM imidazole wash. The target protein was collected and used from L5–L11. The gel was stained with Coomassie Brilliant Blue (CBB).

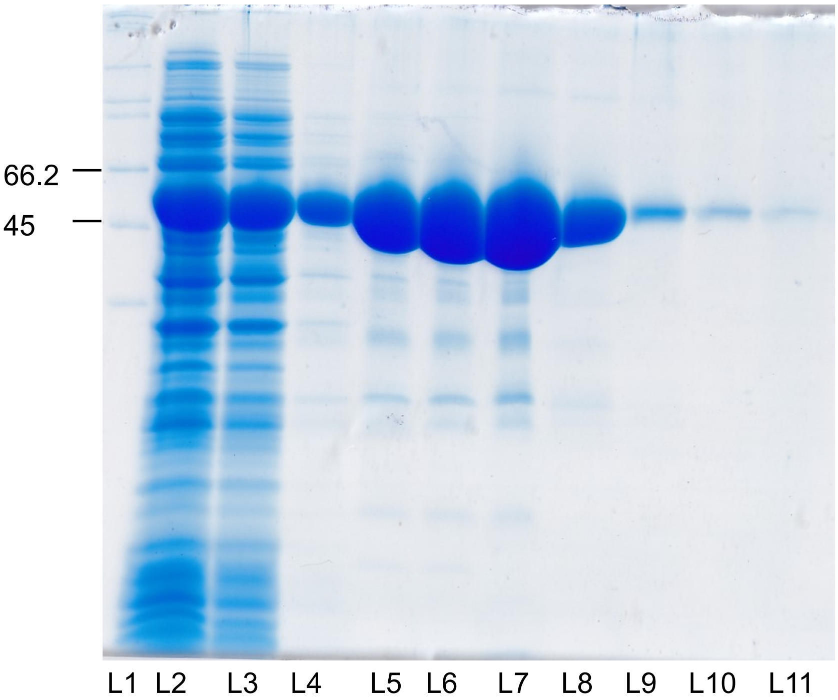


**Supplementary Fig. 8. Full gel depicting the purification of the mutant PchD H503R.** Gel lanes depict fractions taken during Ni-sepharose chromatography (Nacalai Tesque, Inc.) and are as follows: L1 = protein marker, L2 = flow through, L3= wash, L4 = 10 mM imidazole wash, L5 = 20 mM imidazole wash, L6 = 40 mM imidazole wash, L7 = 80 mM imidazole wash, L8 = 100 mM imidazole wash, L9 = 150 mM imidazole wash, L10 = 200 mM imidazole wash, and L11 = 400 mM imidazole wash. The target protein was collected and used from L5–L11. The gel was stained with Coomassie Brilliant Blue (CBB).

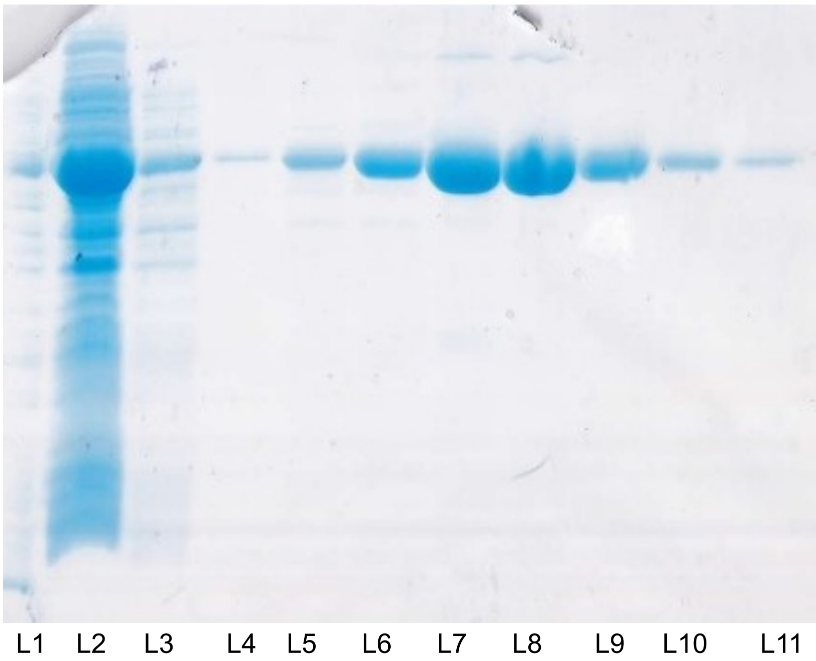


**Supplementary Fig. 9. Full gel depicting the purification of the mutant PchD Q500R/H503R.** Gel lanes depict fractions taken during Ni-sepharose chromatography (Nacalai Tesque, Inc.) and are as follows: L1 = protein marker, L2 = flow through, L3= wash, L4 = 10 mM imidazole wash, L5 = 20 mM imidazole wash, L6 = 40 mM imidazole wash, L7 = 80 mM imidazole wash, L8 = 100 mM imidazole wash, L9 = 150 mM imidazole wash, L10 = 200 mM imidazole wash, and L11 = 400 mM imidazole wash. The target protein was collected and used from L5–L11. The gel was stained with Coomassie Brilliant Blue (CBB).

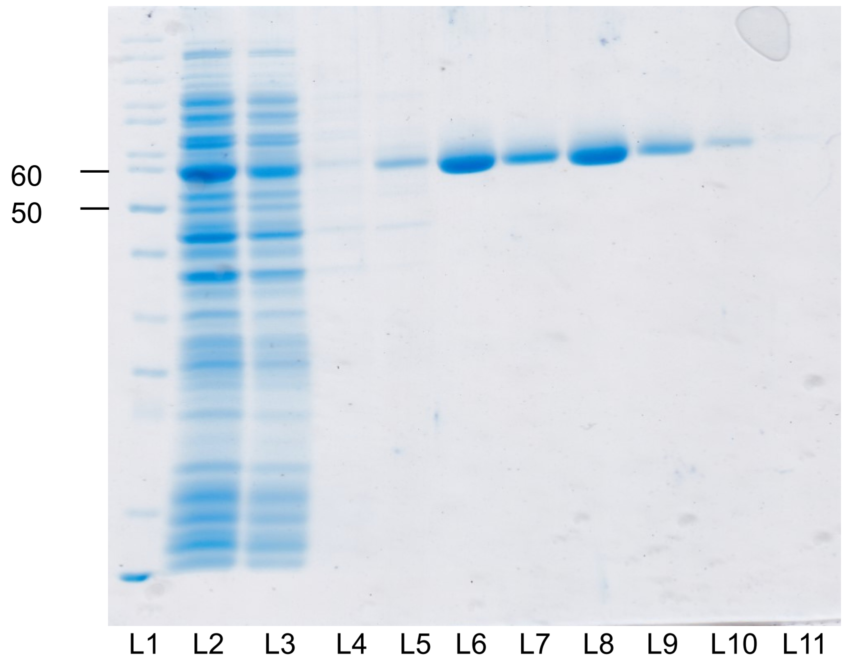


**Supplementary Fig. 10. Full gel depicting the purification of the mutant VibE N239C.** Gel lanes depict fractions taken during Ni-sepharose chromatography (Nacalai Tesque, Inc.) and are as follows: L1 = protein marker, L2 = flow through, L3= wash, L4 = 10 mM imidazole wash, L5 = 20 mM imidazole wash, L6 = 40 mM imidazole wash, L7 = 80 mM imidazole wash, L8 = 100 mM imidazole wash, L9 = 150 mM imidazole wash, L10 = 200 mM imidazole wash, and L11 = 400 mM imidazole wash. The target protein was collected and used from L5–L10. The gel was stained with Coomassie Brilliant Blue (CBB).

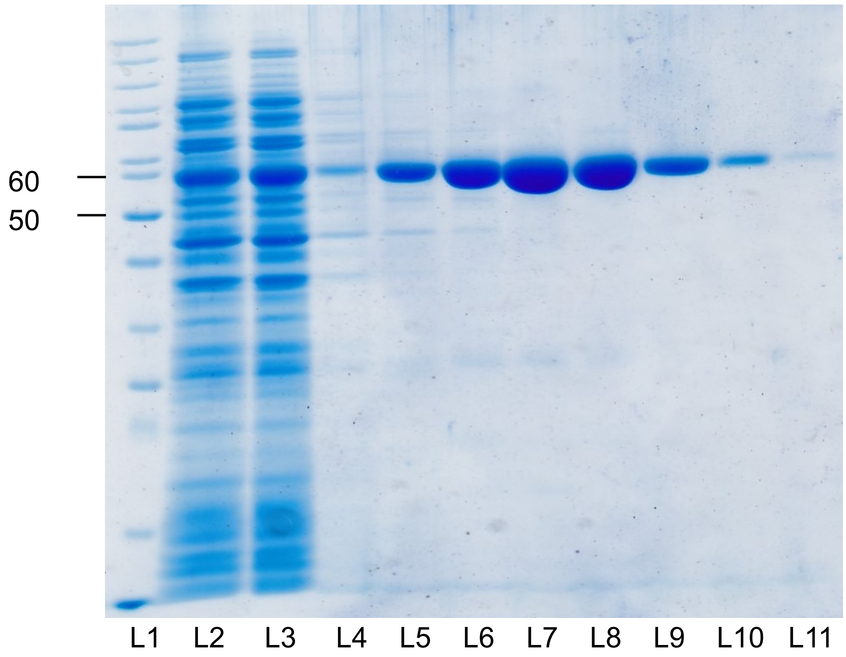


**Supplementary Fig. 11. Full gel depicting the purification of the mutant VibE N239C/H498R.** Gel lanes depict fractions taken during Ni-sepharose chromatography (Nacalai Tesque, Inc.) and are as follows: L1 = protein marker, L2 = flow through, L3= wash, L4 = 10 mM imidazole wash, L5 = 20 mM imidazole wash, L6 = 40 mM imidazole wash, L7 = 80 mM imidazole wash, L8 = 100 mM imidazole wash, L9 = 150 mM imidazole wash, L10 = 200 mM imidazole wash, and L11 = 400 mM imidazole wash. The target protein was collected and used from L5–L11. The gel was stained with Coomassie Brilliant Blue (CBB).

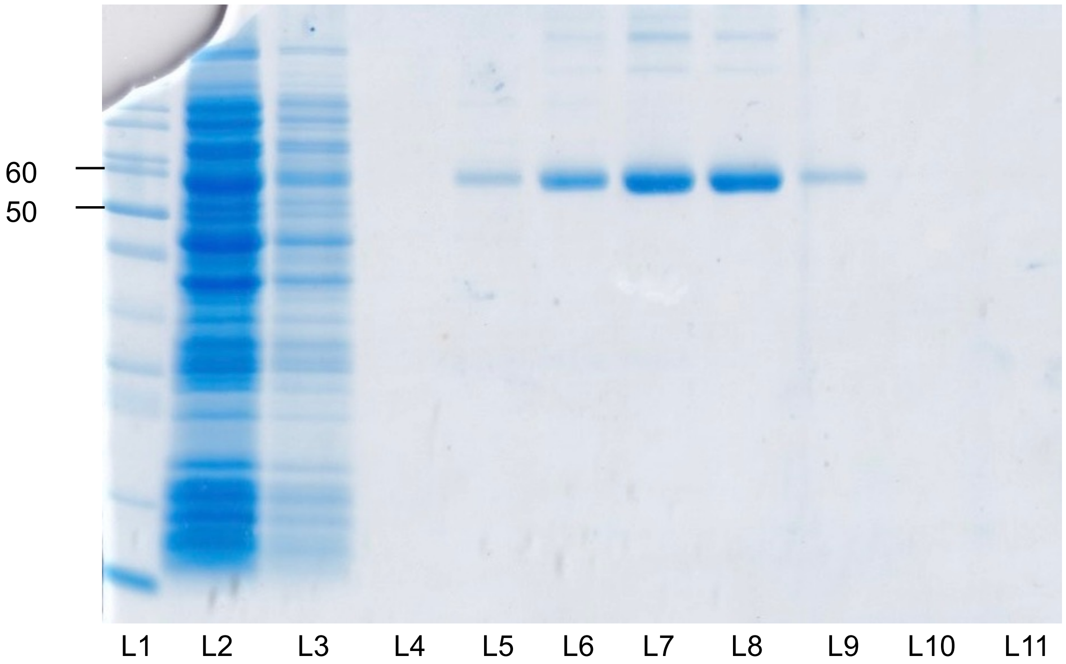


**Supplementary Fig. 12. Full gel depicting the purification of the mutant VibE N239C/Q510R.** Gel lanes depict fractions taken during Ni-sepharose chromatography (Nacalai Tesque, Inc.) and are as follows: L1 = protein marker, L2 = flow through, L3= wash, L4 = 10 mM imidazole wash, L5 = 20 mM imidazole wash, L6 = 40 mM imidazole wash, L7 = 80 mM imidazole wash, L8 = 100 mM imidazole wash, L9 = 150 mM imidazole wash, L10 = 200 mM imidazole wash, and L11 = 400 mM imidazole wash. The target protein was collected and used from L5–L11. The gel was stained with Coomassie Brilliant Blue (CBB).

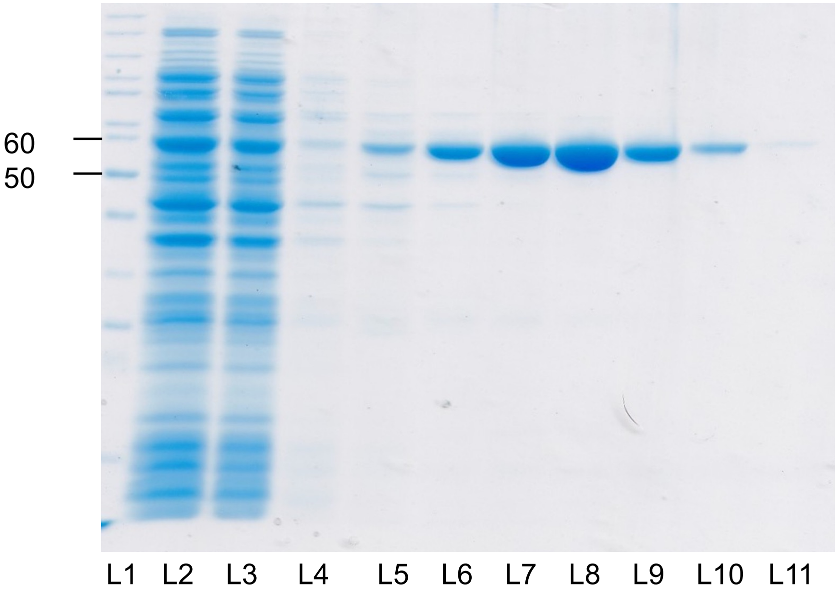


**Supplementary Fig. 13. Full gel depicting the purification of the mutant VibE N239C/H498R/Q510R.** Gel lanes depict fractions taken during Ni-sepharose chromatography (Nacalai Tesque, Inc.) and are as follows: L1 = protein marker, L2 = flow through, L3= wash, L4 = 10 mM imidazole wash, L5 = 20 mM imidazole wash, L6 = 40 mM imidazole wash, L7 = 80 mM imidazole wash, L8 = 100 mM imidazole wash, L9 = 150 mM imidazole wash, L10 = 200 mM imidazole wash, and L11 = 400 mM imidazole wash. The target protein was collected and used from L6–L11. The gel was stained with Coomassie Brilliant Blue (CBB).

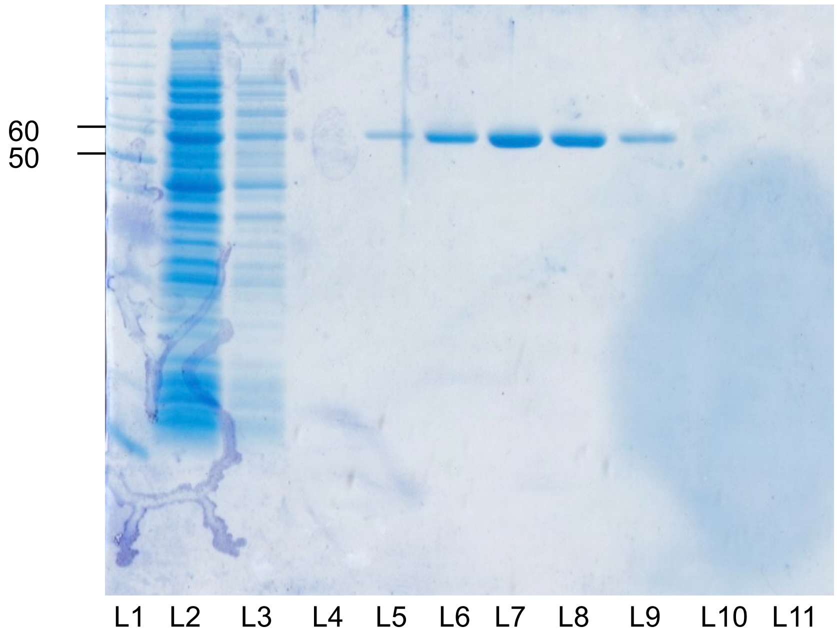


**Supplementary Fig. 14. Full gel depicting the purification of the mutant VibE N239C/K494R/H498R/Q510R.** Gel lanes depict fractions taken during Ni-sepharose chromatography (Nacalai Tesque, Inc.) and are as follows: L1 = protein marker, L2 = flow through, L3= wash, L4 = 10 mM imidazole wash, L5 = 20 mM imidazole wash, L6 = 40 mM imidazole wash, L7 = 80 mM imidazole wash, L8 = 100 mM imidazole wash, L9 = 150 mM imidazole wash, L10 = 200 mM imidazole wash, and L11 = 400 mM imidazole wash. The target protein was collected and used from L5–L9. The gel was stained with Coomassie Brilliant Blue (CBB).

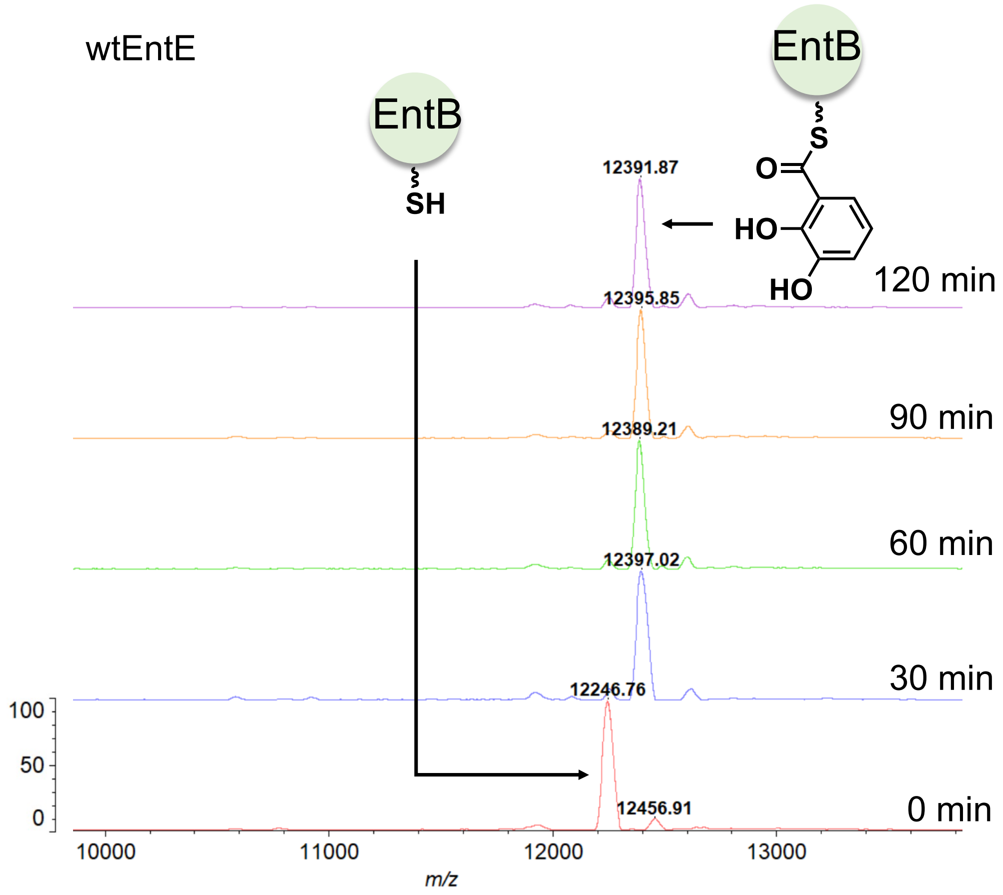


**Supplementary Fig. 15. Expanded views of the MALDI-TOF mass spectra in Fig. 3A.** Time-course MALDI-TOF mass spectra for DHB loading onto EntB(ArCP) by wtEntE. Spectra were acquired at 0, 30, 60, 90, and 120 min. Peaks corresponding to the *holo* and DHB-loaded forms of EntB(ArCP) are indicated.

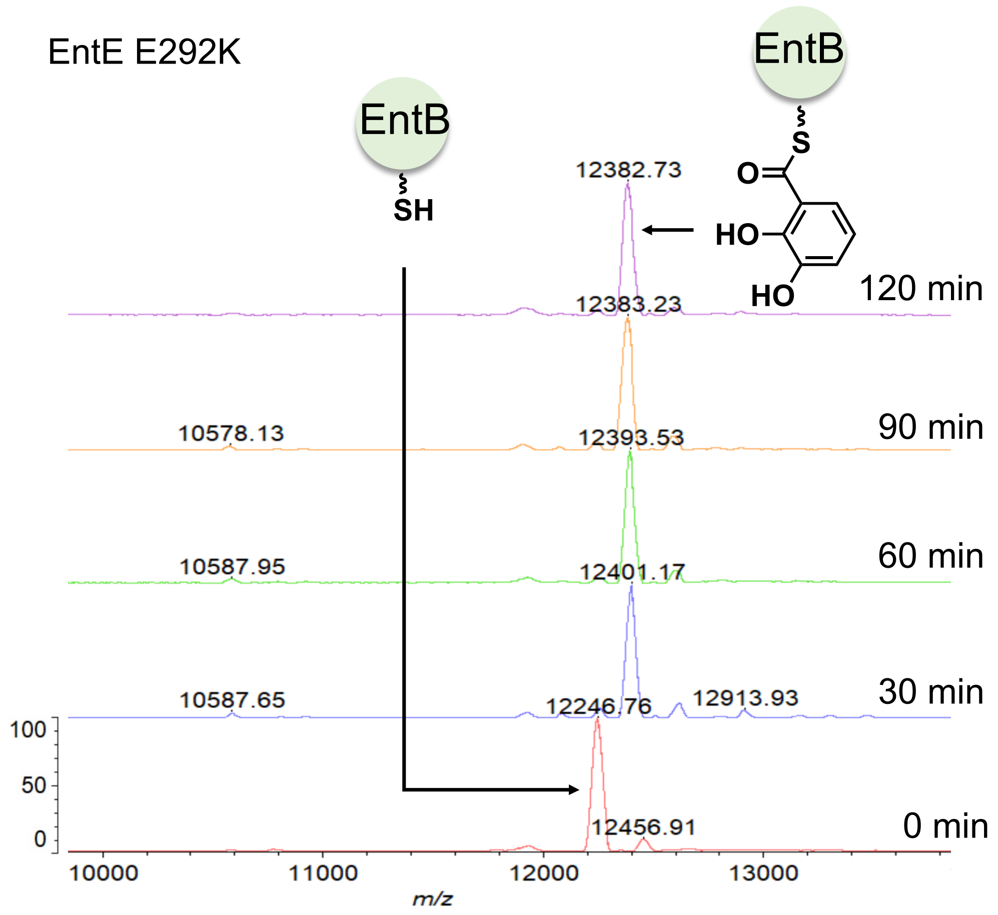


**Supplementary Fig. 16. Expanded views of the MALDI-TOF mass spectra in Fig. 3B.** Time-course MALDI-TOF mass spectra for DHB loading onto EntB(ArCP) by EntE E292K. Spectra were acquired at 0, 30, 60, 90, and 120 min. Peaks corresponding to the *holo* and DHB-loaded forms of EntB(ArCP) are indicated.

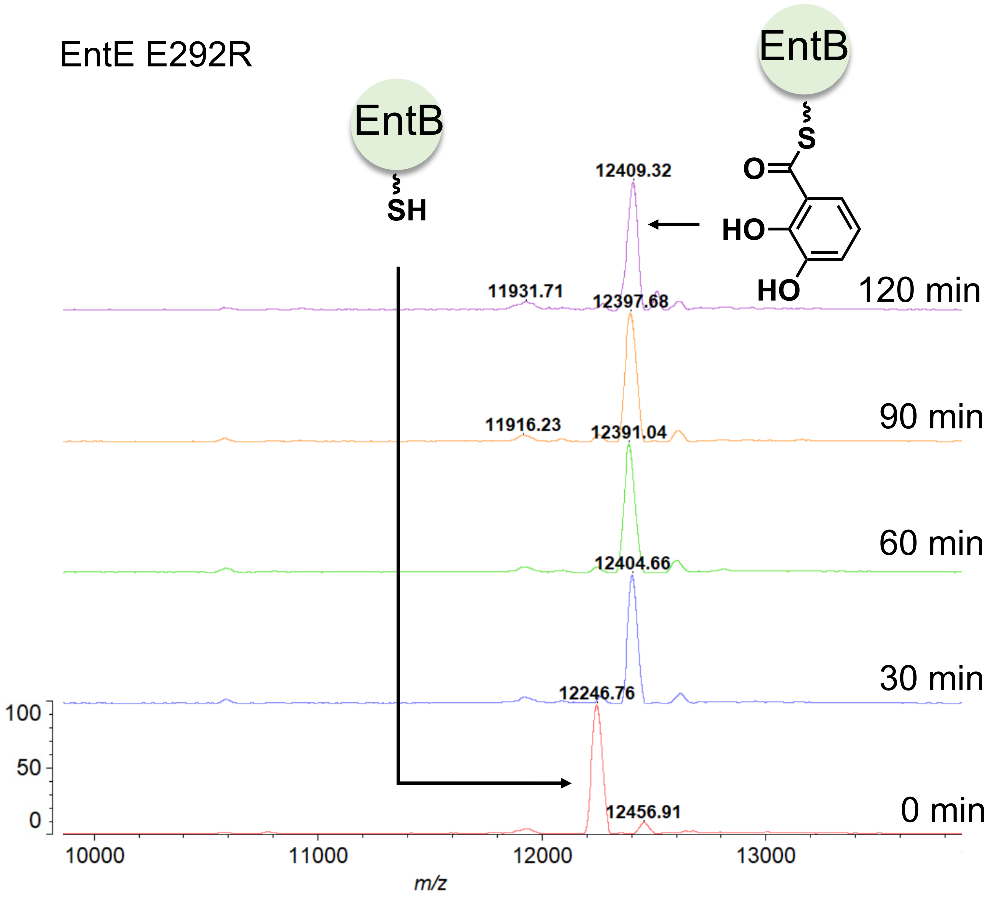


**Supplementary Fig. 17. Expanded views of the MALDI-TOF mass spectra in Fig. 3C.** Time-course MALDI-TOF mass spectra for DHB loading onto EntB(ArCP) by EntE E292R. Spectra were acquired at 0, 30, 60, 90, and 120 min. Peaks corresponding to the *holo* and DHB-loaded forms of EntB(ArCP) are indicated.

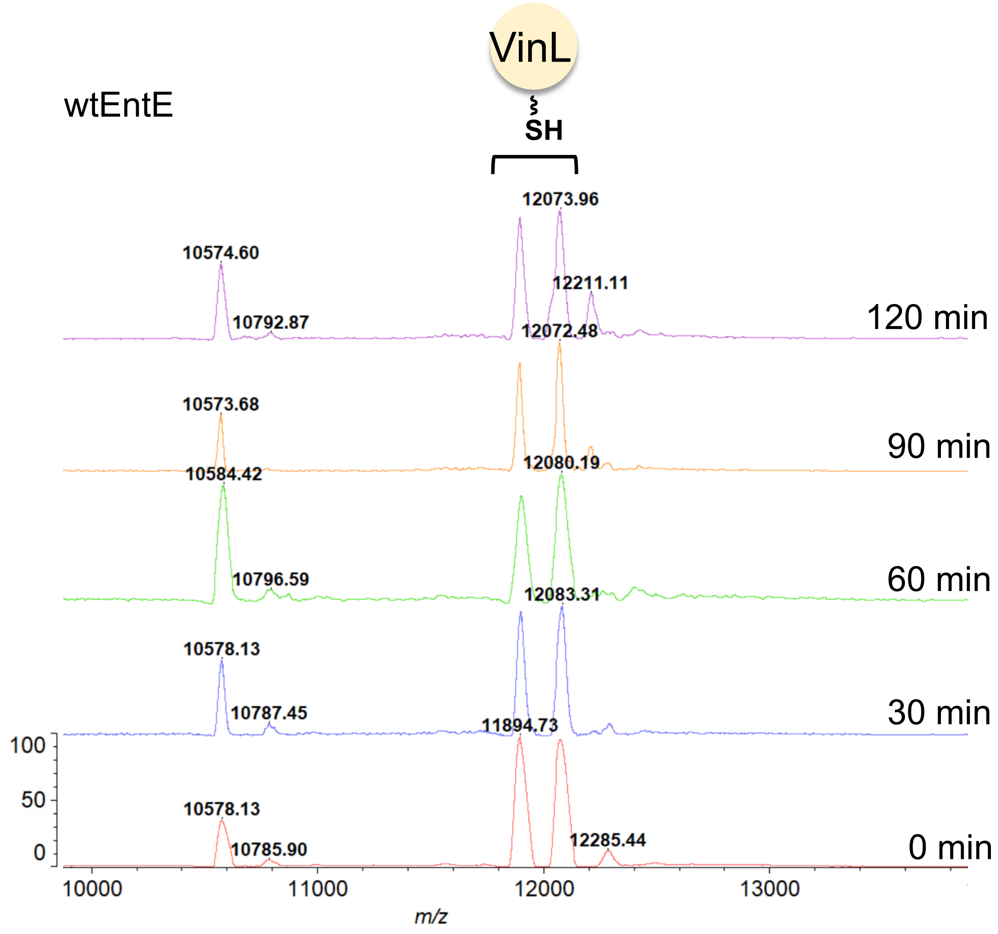


**Supplementary Fig. 18. Expanded views of the MALDI-TOF mass spectra in Fig. 3D.** Time-course MALDI-TOF mass spectra for DHB loading onto VinL by wtEntE. Spectra were acquired at 0, 30, 60, 90, and 120 min. Peaks corresponding to the *holo* and DHB-loaded forms of VinL are indicated.

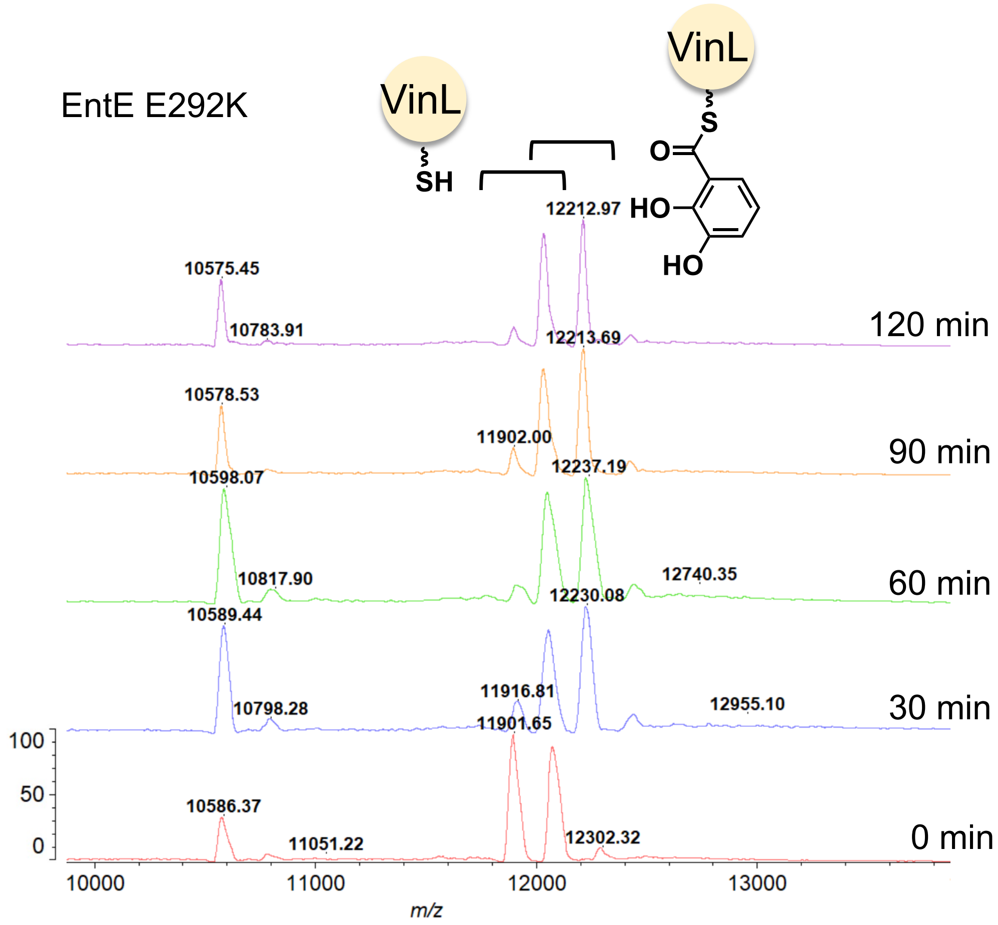


**Supplementary Fig. 19. Expanded views of the MALDI-TOF mass spectra in Fig. 3E.** Time-course MALDI-TOF mass spectra for DHB loading onto VinL by EntE E292K. Spectra were acquired at 0, 30, 60, 90, and 120 min. Peaks corresponding to the *holo* and DHB-loaded forms of VinL are indicated.

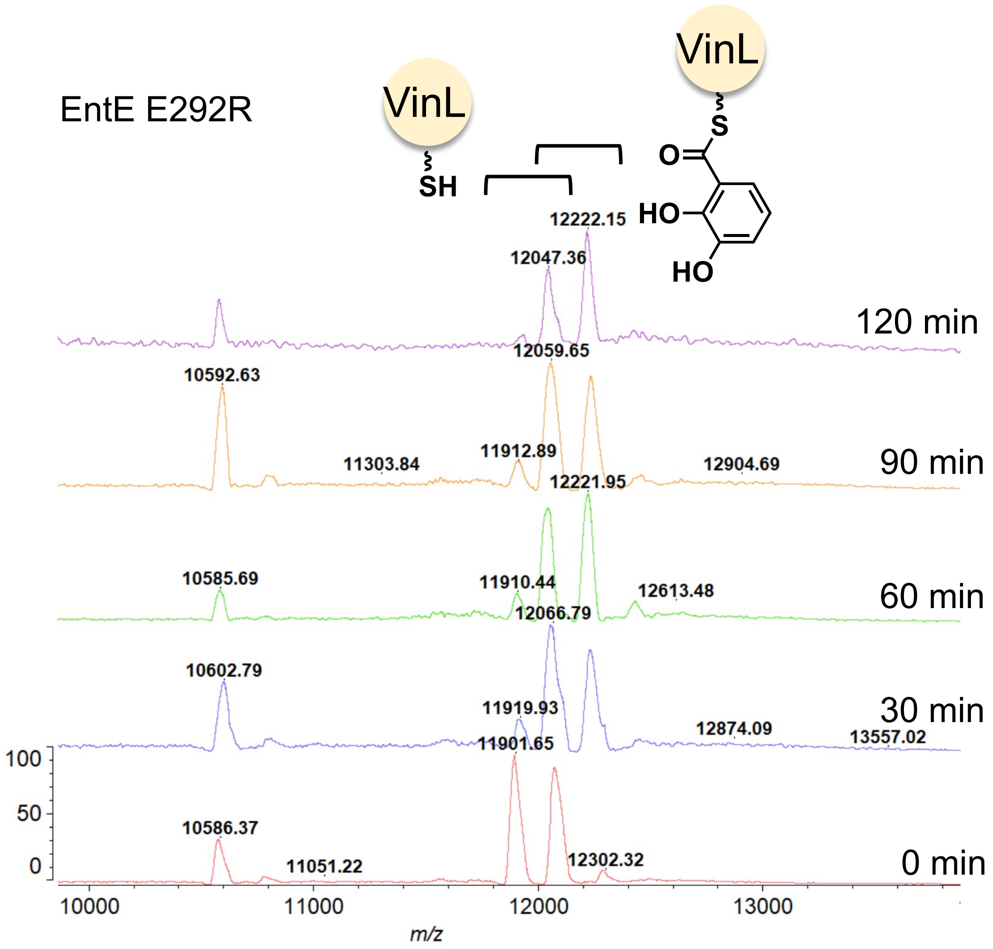


**Supplementary Fig. 20. Expanded views of the MALDI-TOF mass spectra in Fig. 3F.** Time-course MALDI-TOF mass spectra for DHB loading onto VinL by EntE E292R. Spectra were acquired at 0, 30, 60, 90, and 120 min. Peaks corresponding to the *holo* and DHB-loaded forms of VinL are indicated.

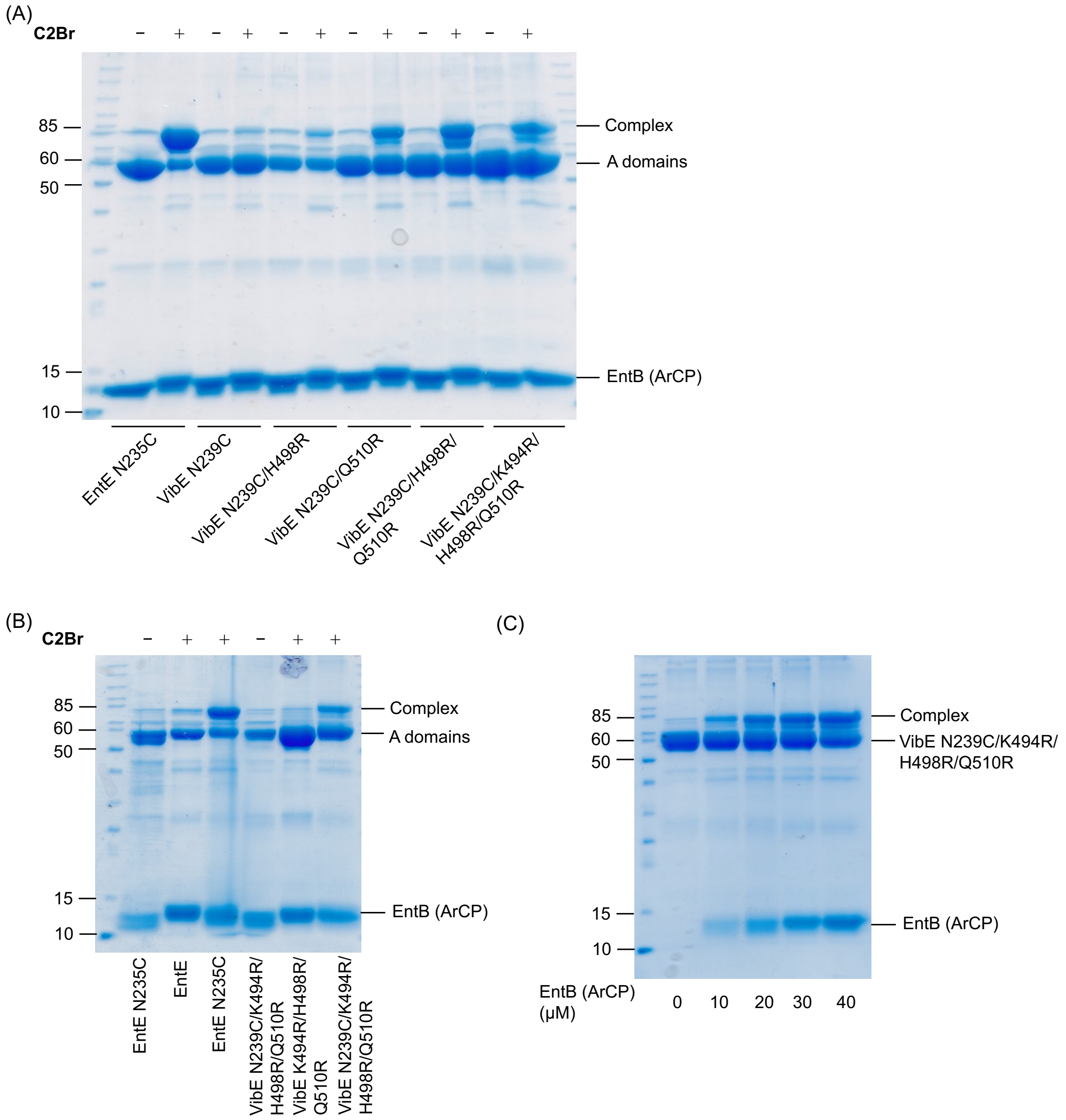


**Supplementary Fig. 21. Uncropped SDS-PAGE gels corresponding to Extended Data Fig. 8.** (A) Extended Data Fig. 8B, crosslinking reactions of EntB(ArCP) with EntE N235C, VibE N239C, VibE N239C/H498R, VibE N239C/Q510R, VibE N239C/H498R/Q510R, and VibE N239C/K494R/H498R/Q510R in the presence or absence of **C2Br**. (B) Extended Data Fig. 8C, crosslinking efficiency and validation of cysteine reactivity. Crosslinking reactions of EntB(ArCP) with wtEntE, EntE N235C, VibE K494R/H498R/Q510R, and VibE N239C/K494R/H498R/Q510R in the presence or absence of **C2Br**. (C) Extended Data Fig. 8D, concentration dependence of crosslinking on EntB(ArCP) concentration. Crosslinking reactions of EntB(ArCP) with VibE N239C/K494R/H498R/Q510R in the presence of **C2Br**. All one-pot crosslinking reactions included treatment with EntB(ArCP), **C2Br**, CoaA, CoaD, CoaE, and Sfp for 1 h at 37 °C. **C2Br**-*crypto*-EntB(ArCP) was then incubated with 5 μM adenylation enzymes for 16 h at 37 °C.

**Supplementary Table 1.** Structural data collection and refinement statistics.

| **VibE-EntB(ArCP) crosslinked complex** | |
|---|---|
| **Data collection** | |
| Beamline | SPring-8 BL45XU |
| Wavelength (Å) | 1.000000 |
| Resolution range (Å) | 48.19–2.38 (2.47–2.38) |
| Space group | $P2_1$ |
| Unit cell parameters | |
| a, b, c (Å) | 76.07, 82.50, 89.21 |
| α, β, γ (°) | 90.000, 93.02, 90.000 |
| Unique reflections | 44,328 (4,641) |
| Multiplicity | 7.3 (9.7) |
| Completeness (%) | 99.9 (100.0) |
| Mean I/σ (I) | 6.9 (2.0) |
| $R_{merge}$ | 0.080 (0.850) |
| $CC_{1/2}$ | 0.997 (0.867) |
| **Refinement** | |
| Resolution range (Å) | 48.19-2.38 |
| $R_{work}$ | 0.228 |
| $R_{free}$ | 0.265 |
| Number of VibE-EntB(ArCP) complex in asymmetric unit | 2 |
| Number of atoms | |
| VibE | 8400 |
| EntB(ArCP) | 989 |
| pantetheine probe | 48 |
| ADP | 54 |
| solvent | 296 |
| Average B-factor (Å$^2$) | |
| VibE | 32.9 |
| EntB(ArCP) | 72.2 |
| pantetheine probe | 44.5 |
| ADP | 30.6 |
| solvent | 27.9 |
| RMS Deviations | |
| bonds lengths (Å) | 0.008 |
| bond angles (°) | 1.781 |
| Ramachandran plot | |
| Favored (%) | 97.3 |
| Allowed (%) | 2.6 |
| Outlier (%) | 0.1 |
| Molprobity Clashscore | 3.82 |
